# Discovery of an Adaptive Neuroimmune Response Driving Itch and Fast Tick Removal with Implications for Preventing Pathogen Transmission

**DOI:** 10.1101/2025.08.22.671835

**Authors:** Johannes S. P. Doehl, Tiago D. Serafim, Serena Doh, Charles S. Grugan, Eva Iniguez, Luana Rogerio, Ronja Frigard, Ranadhir Dey, Pedro Cecilio, Xinglong Gu, Pang-Yen Tseng, Aline Da Silva Moreira, Mahnaz Minai, James Oristian, Hans Ackerman, Steve Brooks, Caroline Percopo, Siu-Ping Ng, Derron A. Alves, Lucas Tirloni, Jennifer M. Anderson, Fabiano Oliveira, Shaden Kamhawi, Daniel E. Sonenshine, Adriana Marques, José M. C. Ribeiro, Mark Hoon, Jesus G. Valenzuela

## Abstract

Acquired tick resistance (ATR) is well characterized in tick-exposed animals, compromising tick fitness through antibody-mediated activation of basophils. Yet, anti-tick vaccines inducing ATR have had limited success. Here, we describe a neuroimmune event preceding ATR that leads to rapid host-mediated tick removal. Tick-sensitized guinea pigs mechanically remove ticks within 3-6 hours via an acquired neuroimmune-induced itch response that correlates with increased dermal expression of itch-associated genes like OSM and skin infiltration by T cells and macrophages, independently of IgG and IgE antibodies. When we expose humans to ticks, a similar immune response is observed. Blocking T cells before tick sensitization prevents immune cell infiltration to bite sites and abrogates scratching and tick removal. This neuroimmune response is independent of Trpv1 as tick-sensitized guinea pigs treated with resiniferatoxin remove ticks effectively. Itch-induced tick removal or IITR offers a novel approach to tick-borne disease prevention through early tick detection and removal.

## INTRODUCTION

Ticks are the most important arthropod vectors of disease in the United States (*1–3*), and they transmit a variety of pathogens including bacteria, viruses and protozoa (*4*). The tick *Ixodes scapularis* (black-legged tick) is considered the most relevant tick species in the United States transmitting human pathogens that cause Lyme disease, human granulocytic anaplasmosis, and babesiosis among other infectious diseases (*3, 4*).

Most tick pathogens are transmitted slowly, requiring between 24 to 48 hours of tick attachment to the host (*5*). This offers a window of opportunity to target the tick before pathogens are transmitted. To date, anti-tick vaccines have focused on identifying tick-derived molecules, mostly salivary or gut proteins, that can prevent tick feeding or kill the tick before pathogens are transmitted (*6*).

The roots of anti-tick vaccines come from the observation that animals exposed multiple times to tick bites become resistant to further tick infestations (*7*). This phenomenon, named acquired tick resistance (ATR), results in reduced tick engorgement, reduced tick egg-masses, premature tick detachment and death of the biting tick (*7–9*). The hallmark of ATR is the presence of basophils at the tick bite site and the requirement for antibodies, and both have been associated with poor tick feeding or compromised tick fitness observed around days 3-4 post attachment (*10, 11*). Attempts to accelerate this response by developing an ATR-mediated anti-tick vaccine that prevents tick feeding and achieves tick detachment by or before 48 hours have only been partially successful, and current vaccines that compromise tick feeding and accelerate tick detachment do so only at 72 hours post tick bite (*6, 12, 13*). Unfortunately, this is too late to prevent transmission of most tick-borne pathogens.

Notably, tick loss at 24-48 hours has been observed in the gold standard model of multiple tick exposures in guinea pigs (GPs) whose mobility was unimpeded (*14*). This strongly suggests that there may be another mechanism, independent of poor tick feeding or loss of tick fitness, that is responsible for the early loss of ticks at 24-48 hours in tick-sensitized animals. A study identified an intriguing correlation between individuals who reported repeated episodes of itch at tick-bite sites and a three-fold reduction in Lyme disease incidence, compared to those that do not feel the itch (*15*). This led us to hypothesize that early tick loss in tick-sensitized animals may be mediated by itch through a rapid host-mediated mechanism that precedes ATR.

Many biting ectoparasites are known to induce itch in afflicted hosts by cutaneous hypersensitivity. Insect bites are thought to mediate itch through a type I hypersensitivity response (*16*), while arachnids, like mites (e.g., scabies), can induce itch through Type I and type IV hypersensitivity responses (*10, 17*). Much is known about itch detection (*18*), including that itch and pain are separate sensations, and that pruritogens are detected by transcriptionally distinct sensory nerves that transmit signals through specialized neural circuits (*19, 20*). In mice, which have been the primary model in the itch-field (*21*), three molecularly distinct populations of sensory neurons, non-peptidergic nociceptors 1 (NP1), NP2, and NP3 (Usoskin nomenclature), are associated with pruritus, with NP3 neurons sensing inflammatory itch (*22*). NP2 and NP3-neurons express the Trpv1 ion-channel together with several respective itch receptors, including histamine receptors. NP2 neurons, which are required for chloroquine-responsiveness, express Mrgpra3, while NP3 neurons express receptors for cytokines, such as the inflammatory mediator Oncostatin M (OSM), which modulates the excitability of these neurons (*23*), as well as IL-31 and the cysteinyl leukotriene receptor (*22*). NP3 neurons also utilize the neurotransmitter Nppb, which activates Npr1-spinal cord neurons (*24*). Many of the key signaling components of NP3 are also present in classes of human DRG-neurons, but differences in expression patterns of some NP3 key genes occur between species (*25–27*). Further research in GPs is still needed to identity the specific neurons for itch (*28*).

Here, we show experimental evidence that bites from *Ixodes scapularis* (black-legged tick) nymphs induce an acquired protective itch/scratch response in *Cavia porcellus* GPs, which results in rapid tick removal to subsequent tick exposures. The host response is acquired 5-6 days after the initial bite and is dependent on an effective adaptive T cell-mediated cellular immune response that triggers a Trpv1-independent itch pathway and a targeted scratch response within hours after biting. The itch/scratch response is localized to the tick bite site and results in the mechanical removal of ticks, hence the term Itch-Induced Tick Removal (IITR). The rapid occurrence of IITR enabling GPs to remove ticks as early as 3 hours after bite, provides opportunities to prevent the transmission of important tick-borne pathogens including *Borrelia, Anaplasma, Ehrlichia, Babesia*, and *Rickettsia* species.

## RESULTS

### Scratching mediates early tick removal without affecting tick fitness

We subjected guinea pig (GPs, *Cavia porcellus*) to a series of weekly-spaced, capsule-free nymphal *Ixodes scapularis* tick exposures to observe frequency of tick loss over the course of 72 hours post tick attachment (fig. S1A). We found that during the second and third tick exposures, GPs lost a median of 53% and 80% of ticks within 48 hours post tick attachment (pta), respectively, compared to only 6% loss for tick-naïve animals during the first tick exposure (Fig. 1A) similarly to what has been previously reported (*7, 29, 30*). To test if this tick loss is due to a direct animal tick removal as suggested by Burke et al. (2005) and not to ATR-driven immunity directly preventing tick feeding (*15*), we used Elizabethan collars on tick-sensitized GPs to prevent animals from reaching the attached ticks (Fig. 1B). Surprisingly, tick loss by sensitized GPs was abrogated by using Elizabethan collars (Fig. 1C), suggesting that ticks are actively removed by the GPs. Of note, at 48 hours and 72 hours pta, tick survival and tick feeding/engorgement success, measured by the tick scutal index, were comparable between the restricted 4-times tick-sensitized and tick-naïve groups (fig. S1, B to E). This indicates that this early tick removal is not due to obstruction of tick feeding. Further supporting this hypothesis, tick removal was reconstituted in tick-sensitized animals only after Elizabethan collars were removed (Fig. 1D). To directly test if tick removal is the result of scratching by tick-sensitized GPs, we used an automated behavior observational research system (EthoVision, Noldus^TM^) to measure scratch activity. Compared to tick-naïve animals, a significantly greater increase in scratch bouts was observed at the site of tick attachment after the third tick exposure, with comparably low levels of scratching activity in animals with no ticks in both tick-sensitized and tick-naïve GPs (Fig. 1E). Furthermore, an increase in the total scratch bouts correlated with a decrease in the number of attached ticks by 24 hours pta (Fig. 1F, S1F). As scratching is a proxy for itching (*31, 32*), our results suggest that itch is an acquired response to ticks, and it is responsible for the early tick removal in tick-sensitized GPs.

**Figure 1.**
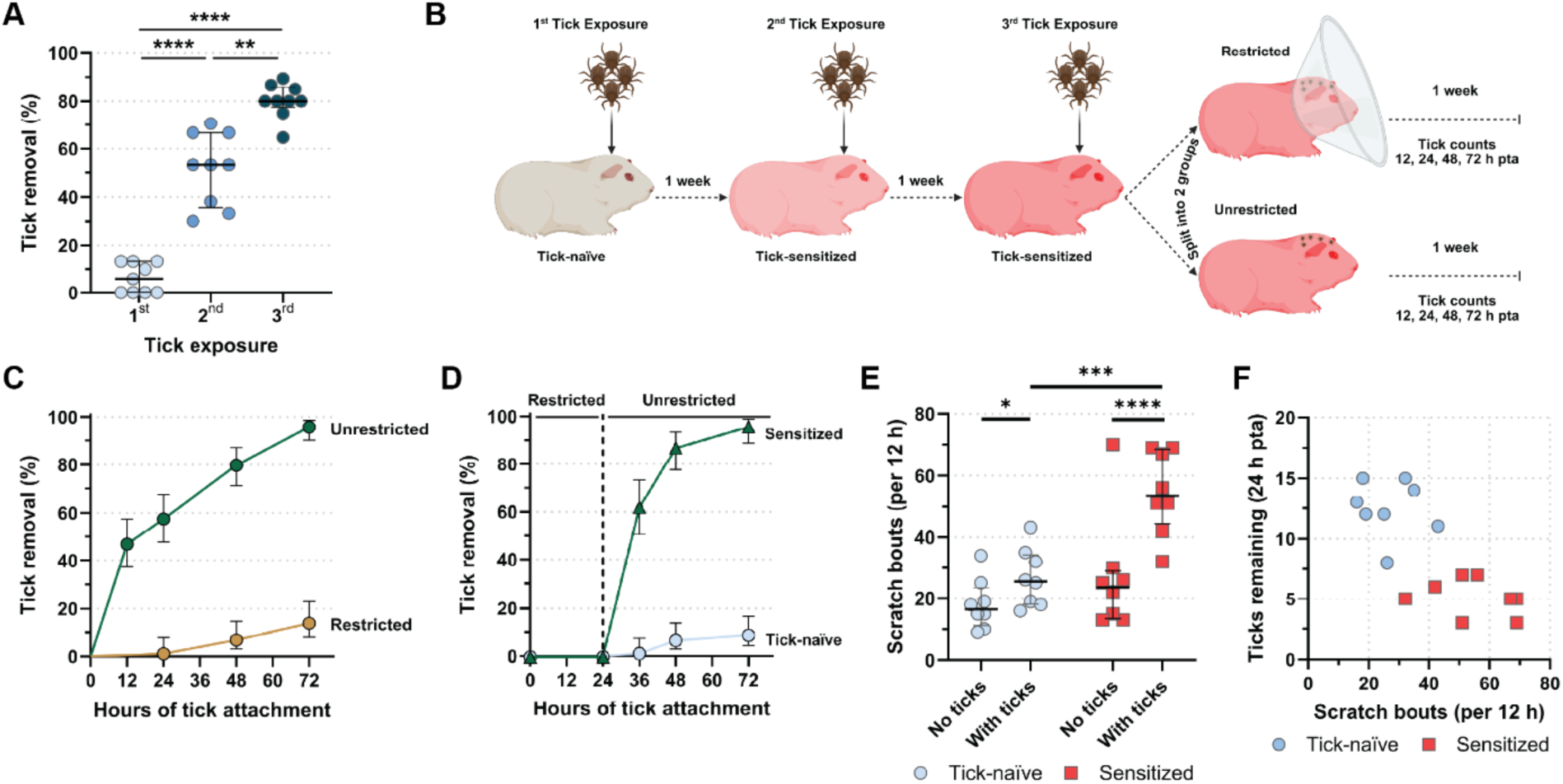
Local scratching mediates rapid tick removal. (**A**) Tick removal in percent of total ticks placed per guinea pig at 48 h post tick attachment (pta) after repeated weekly tick exposures (*N*=15 ticks/guinea pig (GP); *N*=9 GPs). Pairwise estimated marginal means: 1^st^ vs. 2^nd^: p<0.0001; 1^st^ vs. 3^rd^: p<0.0001; 2^nd^ vs. 3^rd^: p=0.004. Median ± IQR shown. (**B**) Experimental concept of **C**. (**C, D**) Smoothed, inverted Kaplan-Meyer plot showing probability of tick removal (%) in (**C**) restricted (*N*=86 ticks) and unrestricted (*N*=94 ticks) tick-access in 5x-exposed GPs (*N*=7/group), and (**D**) tick-naïve (*N*=93 ticks) and 6x-exposed (*N*=68 ticks) GPs (*N*=6/group) after E-collar-removal 24 h pta. (**C,D**) Analysis by Peto & Peto modification of the Gehan-Wilcoxon test: (**C**) p<0.0001, (**D**) p<0.0001. 95% CI is shown. (**E**) Scratch bouts/12 h in tick-naïve and tick-sensitized GPs (*N*=8/group) without, and with attached ticks (*N*=181 ticks and *N*=169 ticks, respectively). Pairwise estimated marginal means: tick-naïve without and with ticks: p=0.0231; tick-sensitized without and with ticks: <0.0001, tick-naïve vs tick-sensitized guinea pigs both with ticks: p<0.0001. Median ± IQR shown. (**F**) Correlation plot of scratch bouts/12 h and ticks remaining (24 h pta) in tick-naïve and tick-sensitized GPs (*N*=8/group). One-way MANOVA: p<0.0001. For detailed statistics see supplementary statistics report.

### Itch-induced tick removal is associated with a skin cellular infiltrate at the bite site that is antibody independent

Given our findings that early tick removal is induced by itch and is an acquired response developed after one tick exposure (priming event), we reasoned that itch may be mediated by an adaptive immune response. Scratching is classically associated with IgE and IgG responses and activation of basophils (*33*) and mast cells (*34*). Surprisingly, anti-tick saliva IgG and total IgE levels in serum remain low in GPs up to week 4 and week 10 pta, respectively, despite a second, third and fourth tick sensitization events on a weekly basis, when IITR responses are fully developed (Fig. 2, A and B), indicating that IITR responses are independent of anti-tick saliva IgG and total IgE antibody levels.

**Figure 2.**
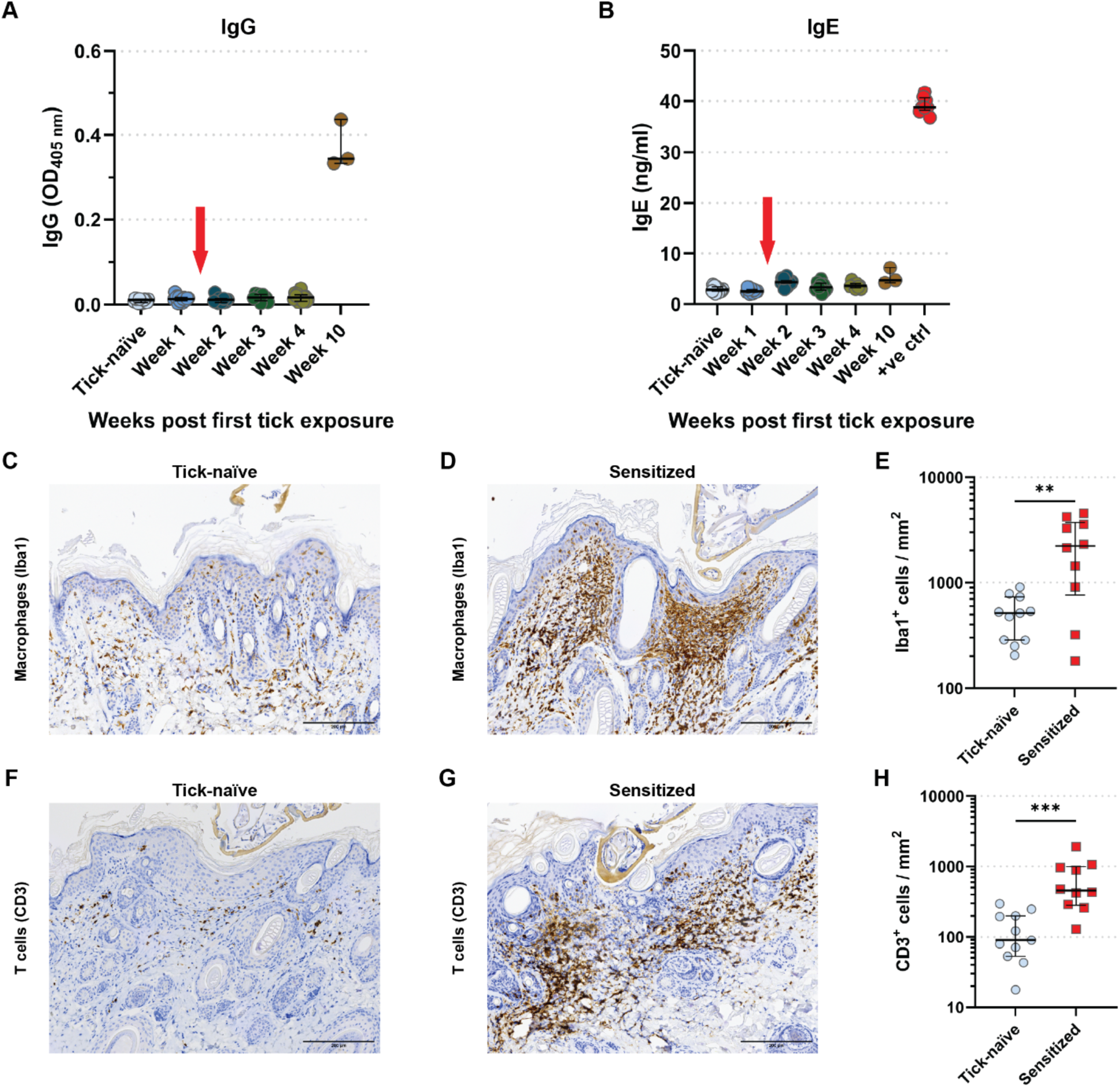
Rapid tick removal is associated with the recruitment of T cells and macrophages but not to IgG or IgE. (**A,B**) ELISA of anti-tick saliva (**A**) IgG and (**B**) total IgE titers in guinea pig (GP) sera pre- and up to 10 weeks post first tick-exposure. Pairwise estimated marginal means: (**A**) All other time-points versus Weeks 10: p<0.0001; (**B**) All other time-points versus positive control (+ve ctrl): p<0.0001. Median ± IQR shown. (**C,D,F,G**) IHC of skin biopsies from tick-bite sites collected at 48 h post tick attachment (pta) from tick-naïve (**C,F)** and tick-sensitized (**D,G**) GPs probed with anti-Iba-1 (macrophages; **C,D)** or anti-CD3 (T cells; **F,G**) antibodies. Scale bar: 200 µm. (**E,H**) Scatter plot: Counts of (**E**) Iba-1^+^ (macrophages), and (**H**) CD3^+^ (T cells) cells in skin sections of biopsies collected at 48 h pta from the 1^st^ exposure of tick-naïve (*N*=11 sections), and the 3^rd^ exposures of tick-sensitized (*N*=10 sections) GPs. Analyzed by (**E**) Welch’s t test: p=0.0002, and (**H**) Mann-Whitney U test: p=0.0058. Median ± IQR shown. For detailed statistics see supplementary report.

We next investigated the recruitment of immune cells to the tick-bite site in the skin of tick-naïve and three-times tick-sensitized GPs at tick bite sites (fig. S2A). At 24 hours pta, tick-sensitized GPs exhibited a strong cellular infiltration (fig. S2A) that was absent in naïve animals (fig. S2B) at the tick-bite site. The skin infiltrate from tick-sensitized GPs consisted of inflammatory cells including Iba-1^+^ macrophages (Fig. 2, C to E) and CD3^+^ T cells (Fig. 2, F to H) that were significantly increased compared to tick-naïve GPs. This cellular infiltration in tick-sensitized animals began as early as 6 hours pta and steadily increased at 12, 24, and 48 hours pta (fig. S3). A few eosinophils were observed in the superficial and mid dermis of tick-sensitized GPs (fig. S2C) and rarely observed in tick-naïve GPs (fig. S2D). The presence of mast cells was at base levels in both tick-sensitized and tick-naïve GPs (fig. S2, A - D).

### Itch-induced tick removal is dependent on the adaptive cellular immune response to tick bites

To directly evaluate whether IITR is dependent on the acquired cellular immune response observed in tick-sensitized GPs, we used the immunosuppressor FTY720. FTY720 prevents the egress of naïve lymphocytes from the lymph nodes, effectively depleting them from circulation and consequently preventing the development of acquired T cell memory immune responses (*35*). GPs were treated daily with FTY720 starting one week prior to the first tick exposure and continuing throughout the first and the second tick exposures (fig. S4A). Consistent with its known effects, we found that FTY720-treated sensitized GPs had a significantly reduced number of circulating lymphocytes compared to sensitized PBS-treated or untreated tick-naïve GPs prior to the first tick exposure (Fig. 3A). Notably, during a second tick exposure, FTY720-treated GPs failed to show significant tick removal while the PBS-treated GPs removed ticks over 48 hours pta (p<0.0001; Fig. 3B). Loss of IITR correlated to an overall reduction of the dermal cellular infiltrate in FTY720-treated (fig. S4B) compared to PBS-treated (fig. S4C) GPs in twice sensitized animals at 48 hours pta. Specifically, treatment with FTY720 significantly decreased the recruitment of Iba-1^+^ macrophages (Fig. 3, C and D, FTY720), and CD3^+^ T cells (Fig. 3, E and F, FTY720) to the tick bite site in comparison to PBS-treated animals (Fig. 3, C to F, PBS). Furthermore, using laser speckle as a proxy for local skin cellular recruitment, we observed that continuous treatment of FTY720 abolished the robust increase in skin blood flow or cell recruitment to the tick bite site (Fig. 3, G and H, FTY720) observed in PBS-treated GPs (Fig. 3, G and H, PBS). As previously observed, the high level of cellular recruitment corresponded to an increase in scratch bouts and in tick removal during the second tick exposure among control PBS-injected animals (Fig. 3,I, PBS 2^nd^). By contrast, FTY720-treated GPs showed fewer scratching bouts upon a second tick exposure corresponding to less effective tick removal (Fig. 3, J, FTY720 2^nd^). These data indicate that IITR relies on a T cell-mediated acquired immune response to ticks that resembles, in kinetics and cell recruitment, a Type IV delayed hypersensitivity response (*36*).

**Figure 3.**
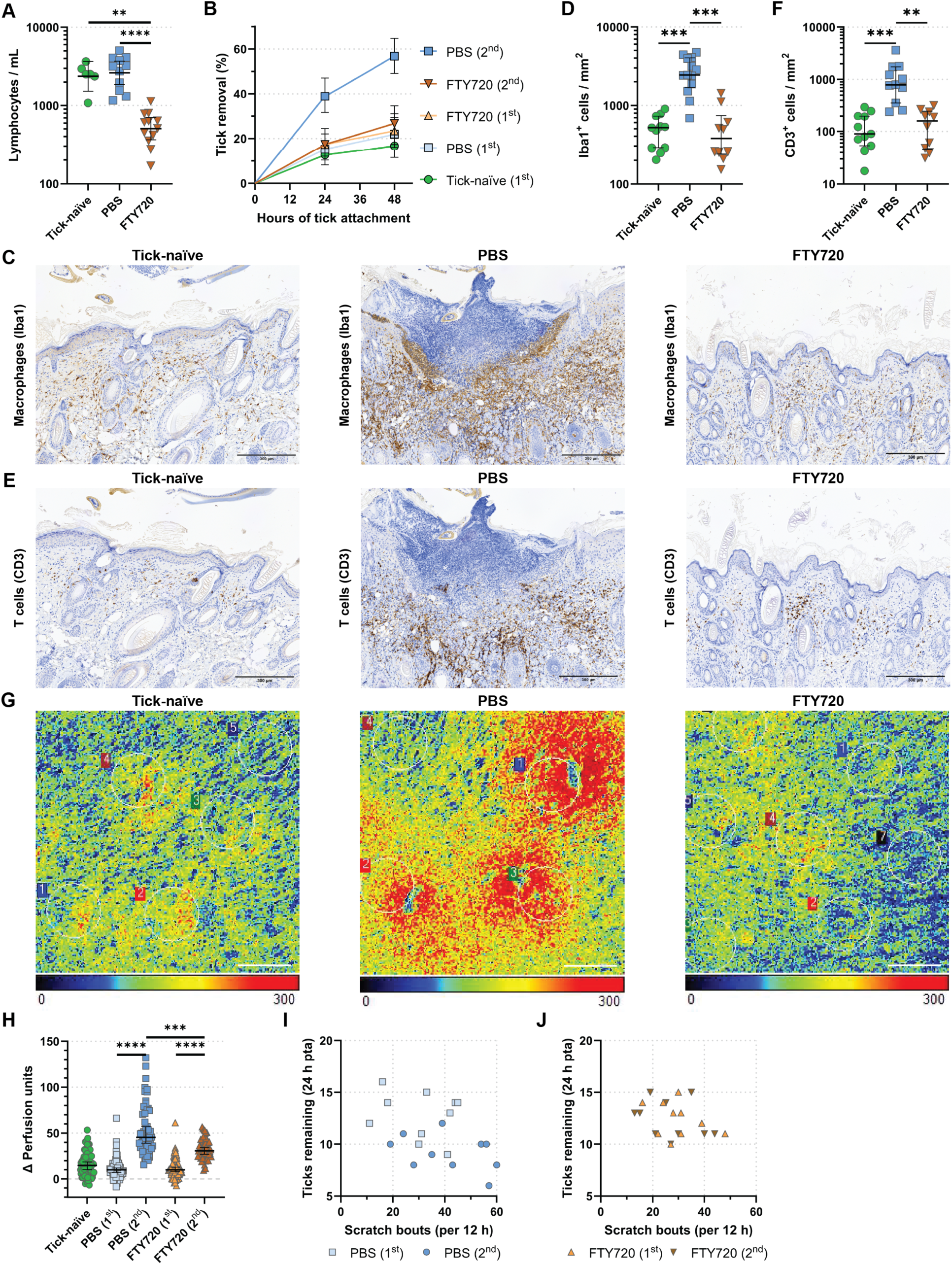
Rapid Tick Removal is dependent on a host adaptive immune response. (**A**) Scatter plots: Lymphocyte counts per milliliter of blood 5 days post FTY720 treatment initiation (tick-naïve: *N*=6 guinea pigs (GPs), PBS: *N*=11 GPs, FTY720: *N*=12 GPs). Analyzed by Dunn’s test: FTY720: Pre-treatment versus Pre-infestation: p=0.0005; Pre-treatment versus post-infestation: p>0.0001. Median ± IQR shown. (**B**) Smoothed, inverted Kaplan-Meyer plot showing probability of tick removal (%) during a 1^st^ exposure of tick-naïve, and a 1^st^ and 2^nd^ exposures of PBS- or FTY720-treated GPs at 24 and 48 h post tick attachment (pta). Analysis by Peto & Peto modification of the Gehan-Wilcoxon test: All groups versus PBS (2^nd^ tick exposure): p<0.0001. 95% CI is shown. *N*=10/group. (**C,E**) IHC of skin biopsies from tick-bite sites collected at 48 h pta of tick-naïve and tick-sensitized (PBS- or FTY720-treated) GPs probed with (**C**) anti-Iba-1 (macrophages) or (**E**) anti-CD3 (T cells) antibodies. Scale bar: 300 µm. (**D,F**) Scatter plots: Counts of (**D**) Iba-1^+^ (macrophages), and (**F**) CD3^+^ (T cells) cells in skin sections of biopsies collected at 48 h pta from the 1^st^ exposure of tick-naïve (*N*=11 sections, either), and the 2^nd^ exposures of PBS- (*N*=13 and *N*=11sections, respectively) or FTY720-treated GPs (*N*=10 sections, either). Analyzed by Dunn’s test: (**D**) 2^nd^ tick exposure of FTY720- versus PBS-treated GPs: p=0.0001, 1^st^ tick exposure of tick-naïve versus 2^nd^ tick exposure of PBS-treated GPs: p=0.0003; and (**F**) 2^nd^ tick exposure of FTY720- versus PBS-treated GPs: p=0.0009, 1^st^ tick exposure of tick-naïve versus 2^nd^ tick exposure of PBS-treated GPs: p=0.0003. Median ± IQR shown. (**G,H**) Laser speckle measurements of tick-bite sites of tick-naïve and tick-sensitized (PBS- or FTY720-treated) GPs. (**G**) Perfusion images. Blue (0) = no signal, red (300) = high signal. Scale bar, 2 mm. (**H**) Scatter plot: Delta Perfusion units (tick-bite site signal - image background signal). Pairwise estimated marginal means: PBS (1^st^ versus 2^nd^ tick exposure): p<0.0001, FTY720 (1^st^ versus 2^nd^ tick exposure): p<0.0001, 2^nd^ tick exposure (FTY720 versus PBS): p=0.006. Median ± IQR shown. (**I, J**) Correlation scatter plot of total scratch bouts/12 h versus total ticks remaining at 24 h pta of 1^st^ and 2^nd^ time exposed GPs (*N*=10/group). (**I**) mock-treated (PBS). (**J**) FTY720-treated. One-way mixed MANOVA: (**I**) p<0.001; (**J**) p=0.069. For detailed statistics see supplementary report.

### Itch-induced tick removal is independent of the Trpv1 pathway

Having established that IITR is evoked by itch, we wanted to identify potential itch signatures in tick-sensitized GPs. To achieve this, we analyzed the differential expression of genes in skin cells from GPs sensitized once or three times to ticks (fig. S5). Principal component analysis distinguished three-times tick-sensitized GPs from both unbitten and once-exposed GPs (fig. S5A). Further, we noted differential expression of genes and an increased expression of molecules known to modulate the itch response (*23*), including defensins, IL13 and OSM, in three times tick-sensitized GPs compared to unbitten and once-exposed GPs (fig. S5B, Table S1).

Next, we wanted to explore whether the molecular neuronal signaling apparatus that triggers an itch response is present in GPs. For this we examined single cell sequencing data from the dorsal root ganglia (DRG) of naïve GPs from a previous study (*28*) for molecules which might be involved in the detection of potential pruritogens from Table S1. Analysis of these data highlighted two transcriptionally distinct molecular populations of Nppa-positive putative itch sensory neurons (Fig. 4A) with enriched expression of the IL-31 cytokine receptor complex for itch, Osmr and IL31Ra (Fig. 4, B and C) (*23, 37*). Nppa is a close family homolog of Nppb and, like Nppb, activates the Npr1 receptor (*38*). Single cell RNAseq data also indicated that these classes of neurons differed from similar murine and human itch neurons because of their lack of expression of Trpv1 as well as other itch specific genes (Fig. 4D) (*25–27*). To validate these unique expression patterns, we performed a multicolor label in situ hybridization (ISH). These studies confirmed that GP DRG contains a class of neurons expressing Nppa and Osmr (Fig. 4E), IL31R (Fig. 4F), but not Trpv1 (Fig. 4G). In addition, we detected expression of the itch receptor Npr1 in neurons in the superficial lamina of the GP spinal cord (Fig. 4H).

**Figure 4.**
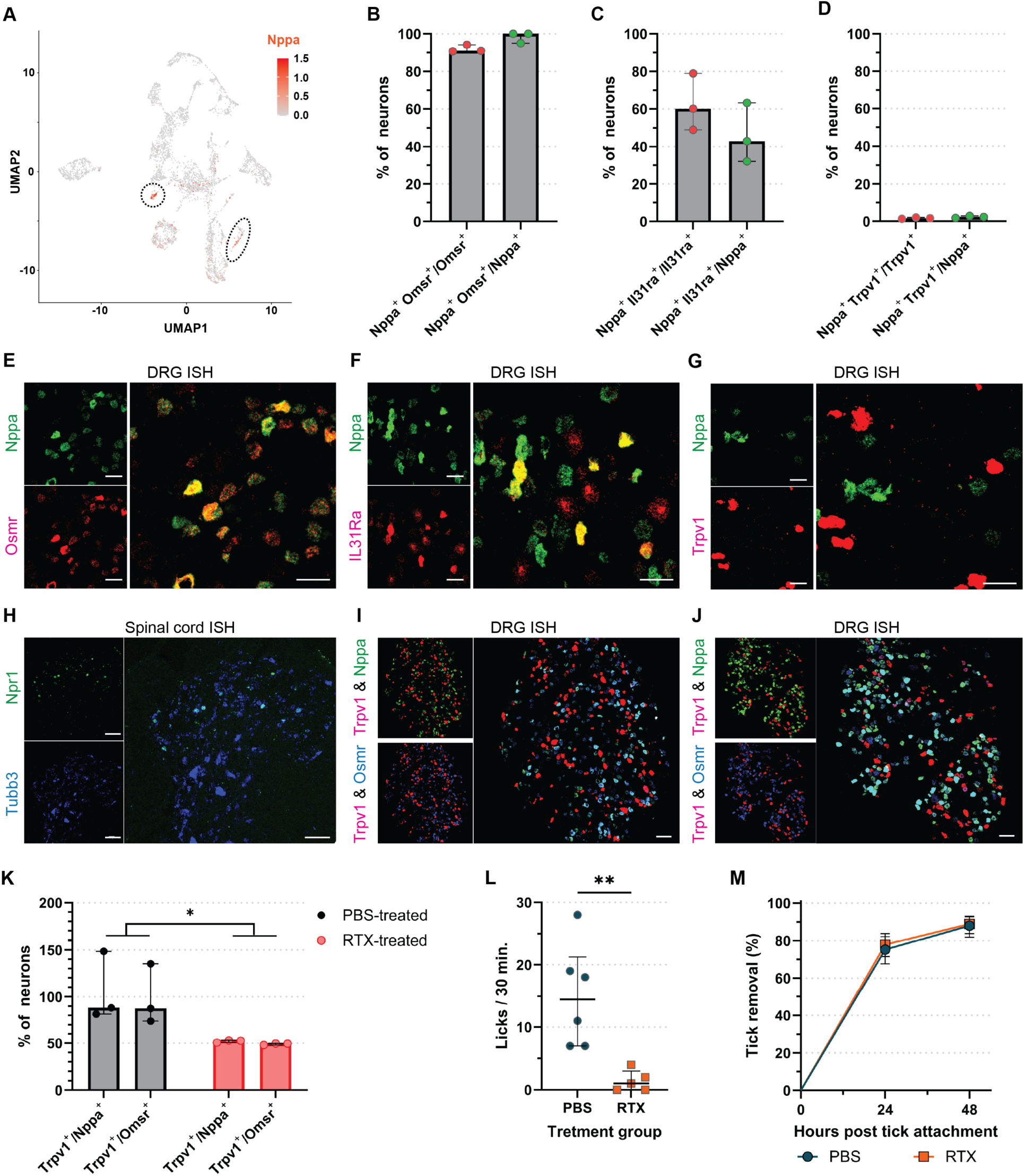
Immune-induced itch in response to tick bite is not mediated through a Trpv1 neural pathway. (**A**) UMAP of scRNA-seq of guinea pig (GP) dorsal root ganglia (DRG) neurons (GEO: GSE201654) highlighting Nppa expression. (**B-D**) Quantification of neurons co-expressing Nppa and (**B**) Osmr, (**C**) IL31Ra, or (**D**) Trpv1 (*N*=3 GPs/staining). (**B-D**) Mann-Whitney U test: (**B**) p=0.1, (**C**) p=0.4, (**D**) p=0.2. Median ± IQR shown. (**E-G**) A representative multiplex ISH image showing co-expression of (**E**) Osmr, (**F**) IL31Ra, or (**G**) Trpv1 on GP DRG Nppa^+^ neurons. Scale bar: 50 µm. (**H**) Expression of Npr1 on Tubb3^+^ GP spinal cord dorsal horn neurons. Scale bar: 50 µm. (**I,J**) A representative multiplex ISH image showing co-expression of Trpv1, Nppa and Osmr on GP DRG neurons in (**I**) PBS-treated (*N*=6) and (**J**) RTX-treated (*N*=5) GPs. Partial ablation of Trpv1 is observed in RTX-treated GPs. Scale bar: 100 µm. (**K**) Quantification of partial Trpv1 ablation in DRG neurons by RTX-treatment. Two-way ANOVA: PBS- versus RTX-treatment: p=0.235. Median ± IQR shown. (**L**) Number of licks to the capsaicin injection site in PBS- and RTX-treated GPs over the course of 30 min. Mann-Whitney U test: p=0.0022. Median ± IQR shown. (**M**) Tick removal on PBS- and RTX-treated GPs. Analysis by Peto & Peto modification of the Gehan-Wilcoxon test: p=0.5381. 95% CI is shown. For detailed statistics see supplementary report.

Given the co-expression of Nppa with itch receptors Osmr and IL31R, and the upregulation of OSM in sensitized GPs, it may be possible that this class of neurons acts through the Npr1 receptor to induce the observed itch-induced tick removal (IITR) response to tick bites. Trpv1 is expressed on GP neurons not expressing Nppa (Fig. 4G). Therefore, we hypothesized that ablation of Trpv1 afferent fibers should not affect the scratch response and consequently tick removal. To test this hypothesis, we treated tick-sensitized GPs with resiniferatoxin (RTX) a Trpv1 agonist which ablates Trpv1-expresing neurons (*39*), or with control vehicle. GPs treated with RTX displayed a large reduction of Trpv1-expressing neurons relative to Nppa and Osmr-neurons (Fig. 4K) and they became insensitive to capsaicin, a Trpv1 agonist (Fig. 4L). Importantly, RTX-treated GPs that are insensitive to capsaicin removed ticks as effectively as control-treated GPs that are sensitive to capsaicin (Fig. 4M). This indicates that tick removal or IITR is independent of the Trpv1 pathway and further supports its potential mediation by Nppa neurons. This contrasts with previous work showing that tick-derived proteins induce itch in naïve mice through the Trpv1 pathway (*40*).

### IITR develops within 5 days after initial tick attachment

To investigate how fast the neuroimmune response leading to IITR develops, we exposed tick-naïve GPs to ticks and collected skin punch biopsies at days 1-5 post-initial tick attachment (Fig. 5, A to E). On day 1 pta, a mild superficial dermal edema was observed (Fig. 5A). On days 2-4, the epidermis at and adjacent to the site of the attached tick was 2-3 times the normal thickness with a minimal dermal inflammatory infiltrate at the tick bite site (Fig. 5, B to D). However, by day 5 pta, we observed a significant increase in dermal cell infiltrate surrounding and separating adnexal structures that extended to the overlying moderately hyperplastic epidermis (Fig. 5E). Cell counts provided further evidence of a significant and sudden increase in cell infiltration at the tick-bite site on day 5 pta (Fig. 5F).

**Figure 5.**
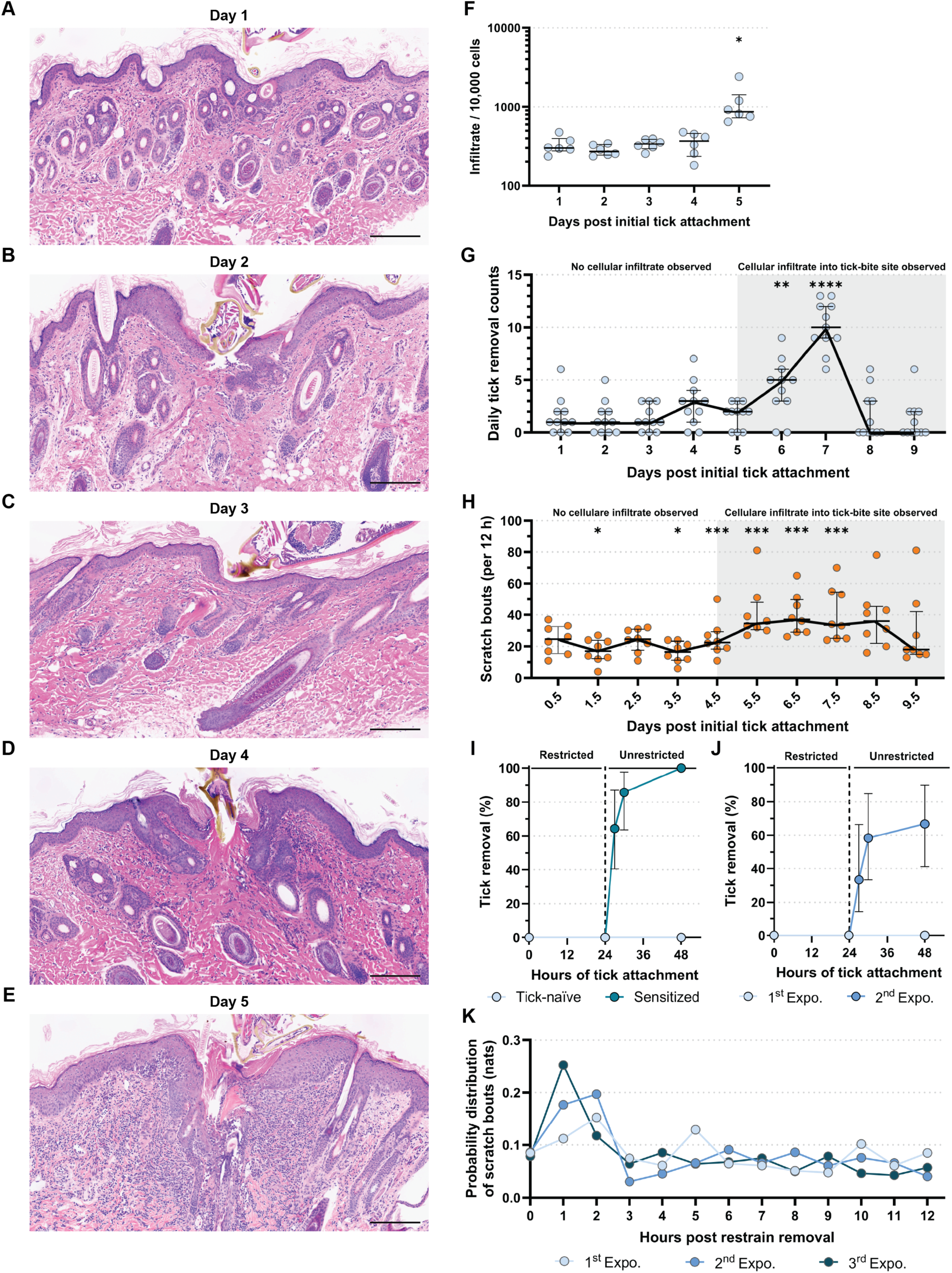
Itch-induced tick removal is rapidly acquired and highly sensitive. (**A-E**) H&E skin cross-sections (>|5µm|<) of tick-bite sites at (**A**) 1 day, (**B**) 2 days, (**C**) 3 days, (**D**) 4 days, and (**E**) 5 days. Scale bar: 200 µm. (**F**) Cellular infiltrate quantification over days 1 – 5 post tick placement (pta). Dunn’s test: Day 1 versus Day 5: p=0.023; Day 2 versus Day 5: p=0.0016. Median ± IQR shown. (**G,H**) Tick-naïve guinea pigs (GPs) continuously exposed to ticks over the course of 10 days. (**G**) Scatter plots showing daily counts of removed ticks. Pairwise estimat d marginal means: 24 h pta versus 144 h pta: p=0.0025; 24 h pta versus 168 h pta: p<0.0001. Median ± IQR shown. (**H**) Scatter plots showing the total scratch bouts per 12 h recording over 10 days. Pairwise estimated marginal means: Day 1 versus day2: p=0.0326; Day 1 versus Day 4: p=0.0184; Day 1 versus Day 6; p=0.0011; Day 1 versus Day 7: p=0.0014; Day 1 versus Day 8: p=0.0017; Day 1 versus Day 9: p=0.0066. Median ± IQR shown. (**I,J**) Smoothed, inverted Kaplan-Meyer plot showing probability of tick removal (%) of a single tick at 27, 30, and 48 h pta after Elizabethan collar removal at 24 h pta. Showing 95% CI. (**I**) Challenge of tick- naïve (*N*=12) and tick-sensitized (*N*=14; 3x sensitized with 15 ticks) GPs. Analysis by Peto & Peto modification of the Gehan-Wilcoxon test: p<0.0001. (**J**) Tick sensitization with single ticks: 1^st^ (*N*=12) and 2^nd^ (*N*=12) tick exposures. Elizabethan collar removal at 24 h pta. Analysis by Peto & Peto modification of the Gehan-Wilcoxon test: p=0.0008. (**K**) Probability distribution (normalization that allows direct comparison of distributions) of hourly scratch bouts measured in natural units of information (nats) against a single attached tick in a 1^st^, 2^nd^, and 3^rd^ tick exposure. Pairwise two-way Kolmogorov-Smirnov test: 1^st^ versus 2^nd^ exposure: p=0.9991; 1^st^ versus 3^rd^ exposure: p=0.9284; 2^nd^ versus 3^rd^ exposure: p=0.9986. For detailed statistics see supplementary report.

To investigate whether the observed sudden increase in cell infiltration 5 days pta corresponds to active tick removal by GPs, we implemented a continuous tick exposure protocol, replacing ticks every 3 days (fig. S6 A) to circumvent natural loss of ticks that fed to repletion (*41, 42*). This allowed us to observe whether increases in targeted scratch bouts and tick removal occurs in conjunction with an increase in the local adaptive cellular immune infiltration. We counted the ticks daily and assessed how many were lost. We found that daily tick removal increased significantly on day 6 and peaked by day 7 pta (Fig. 5G). Most ticks were removed by day 8 pta (Fig. 5 G, fig. S6 B). The observed increase in the rate of tick removal was concurrent with a higher density of cellular infiltrate on day 5 (Fig. 5 E and F), suggesting an association between the cellular infiltrate and induction of IITR. In line with this observation, the increased rate of tick removal also corresponded to the onset of increased scratching at the tick bite site on day 5.5 pta that was maintained up to day 8.5 pta before subsiding on day 9.5 pta when most of the ticks have been removed (Fig. 5, G and H, fig S6 B). Collectively, these data show that the IITR-associated neuroimmune response develops rapidly, within 5 days after a first exposure to tick bites, on GPs not previously exposed to tick bites, leading to an active itch response that results in IITR by 6 days post initial tick exposure.

### Exposure to a single tick bite is sufficient to generate itch-induced tick removal

Next, we sought to determine whether a single tick would elicit IITR in tick-sensitized GPs. We found that single ticks were removed by tick-sensitized GPs within 3-6 hours after collar removal, while tick-naïve GPs failed to remove ticks (Fig. 5I). Remarkably, prior exposure to a single tick was sufficient to acquire the IITR response, efficiently triggering it during a subsequent challenge with one tick (Fig. 5J). This provides evidence that a single tick can prime and sustain effective IITR. In fact, weekly exposures to a single tick over three weeks were as effective in sensitizing GPs as two exposures to 10 ticks (fig. S7A). Importantly, scratch bout frequencies to one tick peaked within the first hours in GPs exposed twice or three times to single ticks and swiftly returned to baseline after tick removal (Fig. 5K), supporting the hypothesis that early tick removal responses are mediated by IITR.

Of note, IITR responses were consistent regardless of whether there was a 1- or 3-week gap between tick exposures (fig. S7B), whether ticks fed for 24 or 72 hours (fig. S7C), or whether male or female GPs were used (fig. S7D). Together, these findings indicate that the elicitation of IITR is highly efficient.

### Itch-induced tick removal is systemic and long-lasting

Next, we investigated whether IITR is a local or systemic response. GPs were exposed three times to 15 ticks at a specific body location and IITR was measured at the third exposure to determine whether the site of tick bites influences its development (fig. S8A). Supporting a systemic response, tick removal efficiency was comparable regardless of the sensitization site (fig. S8, B and C). Importantly, IITR remained intact for at least 6 months after a tick exposure event after one or two tick exposures (fig. S9, A and B), albeit with a slightly attenuated magnitude in the former (fig. S9A). Pertinently, IITR intensity was restored after a new tick exposure, even after a 6-month-long latency (fig. S9C). Cumulatively, our data strongly suggest that IITR is a long-lasting and systemic defense mechanism, allowing a sensitized host to rapidly detect and remove attached ticks.

### Humans experimentally exposed to ticks develop an acquired skin cellular immune response at the bite site comparable to the one observed in guinea pigs

There is evidence that humans develop an itch response to tick bites (*15, 43*). Our results in GPs show that itch-mediated IITR is linked to a skin cellular immune response at the tick bite site. To assess the relevance of these findings in humans, we tested whether volunteers experimentally exposed to bites of *I. scapularis* ticks, the same species we used on GPs, would also develop an acquired skin cellular immune response. Four human volunteers with no history of *I. scapularis* bites were exposed up to three times to 20 *I. scapularis* larval ticks for 24 to 48 hours. Skin biopsies from the 4 volunteers did not show significant cell recruitment at 24 hours after first tick exposure (Fig. 6, A to D), similar to our observations with tick-naïve GPs (fig. S2B). In contrast, after a second or third tick exposures, there was an increase in cell recruitment at the tick-bite site (Fig. 6, E to H) that was significantly higher compared to the response after the first tick exposure (Fig. 6I). Overall, our results indicate that humans mount a rapid acquired skin cellular immune response to tick bites which is central to IITR.

**Figure 6.**
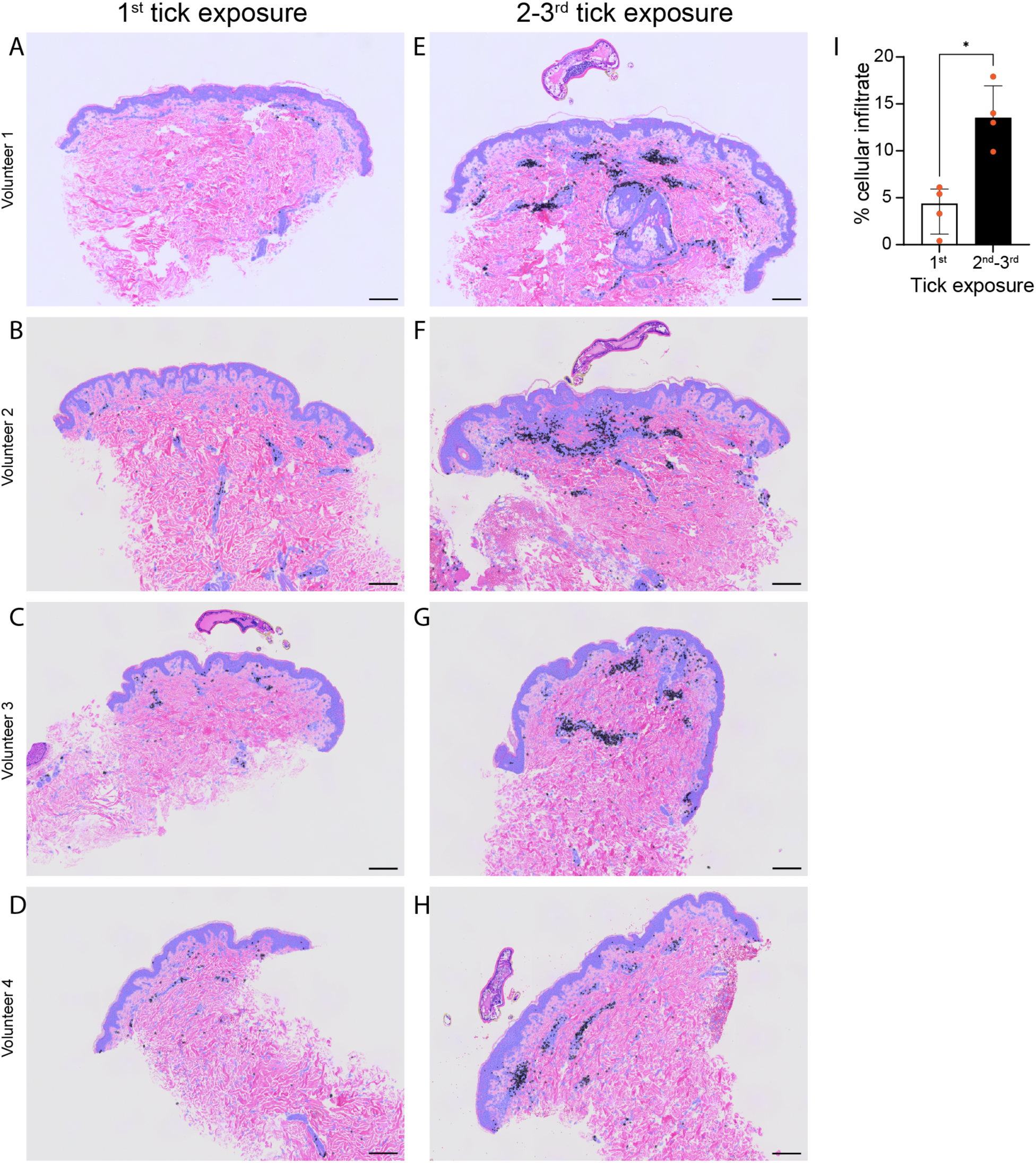
Humans exhibit an adaptive skin cellular immune response at the tick bite site. (A-H) H&E skin biopsy cross-sections (>|5µm|<) of the tick bite site from four human volunteers. (**A-D**) After a 1^st^ tick exposure to 20 *I. scapularis* larval ticks. (**E-H**) After a 2^nd^ or 3^rd^ tick exposure to 10 *I. scapularis* larval ticks. Scale bar: 200 µm. (**I**) Proportion of all cells in the skin biopsies sections (**A-H**) attributed to cellular infiltrate due to larval tick bites comparing 1^st^ tick exposure (A-D) to 2^nd^/3^rd^ tick exposures (**E-H**). Mann-Whitney U test: p=0.0286. Median ± IQR shown. For detailed statistics see supplementary report.

## DISCUSSION

Our findings establish that an acquired cellular immune response in conjunction with activation of neuronal itch pathways are responsible for IITR, an overlooked mechanism observed in tick-sensitized GPs. Itch as a mechanism of tick loss has been proposed previously but never demonstrated or tested (*15, 44*). This work provides a formal demonstration that an immune-induced itch is responsible for a previously overlooked early and active tick removal by tick-sensitized GPs that is independent of IgE or IgG antibodies. The unchanged number of mast cells in the skin of tick-naïve or tick-sensitized GPs suggests that they are also not involved in this itch response. Specifically, IITR is established well before humoral responses develop, with initial tick sensitization of naïve animals taking as little as 5 days. This observed adaptive period, albeit short, indicates that early tick removal is not initiated by innate immune responses alone. Instead, our results suggest that IITR-mediated early tick removal relies on a type IV hypersensitivity response primarily driven by mature T cells and macrophages, that triggers a Trpv1-independent itch pathway leading to localized scratching. It is worth noting that IITR does not exclude the development of ATR and the involvement of IgE- and IgG-activated innate cells at later time periods (*44*). In fact, IITR as described here differs from the previously described ATR in its kinetics, type of immune response, and effect on tick fitness. ATR, unlike IITR, is dependent on antibodies, basophils and mast cells, and compromises tick feeding, tick attachment, and reduces tick weight and egg laying (*44, 45*). These temporal and mechanistic differences clearly distinguish IIRT from ATR, and demonstrate the occurrence of two distinct mechanisms which lead to different anti-ectoparasite responses: an early acquired cell-mediated immunity discovered here that drives rapid itching and active removal of ticks in 3-6 hours, and a later antibody-mediated basophilic immune response that primarily affects ticks fitness (*44, 45*).

The rapid removal of ticks by tick-sensitized animals in as little as 3-6 hours is due to the neuroimmune response described in this work. The activation of neural itch pathways, most likely induced by the observed cell infiltrate, causes the tick-sensitized animal to feel and remove ticks rapidly, in a window of time at or earlier than 24 hours, before infected ticks transmit several major pathogens including *Borrelia* (*5*). This window of time may be vital for development of effective therapeutic strategies for prevention or reduction in the severity of ectoparasite-borne diseases such as Lyme disease, rickettsioses, ehrlichiosis, anaplasmosis, and other infections transmitted by *Ixodes scapularis* and other tick species. These innovative strategies are based on triggering the body’s own itch-based defense systems. In one study, subjects reporting itch in association with tick bites had a lower probability of acquiring Lyme disease (*15*). Based on our findings, we suggest that these individuals probably removed the attached ticks before pathogen injection.

Here we show that GPs naturally induce a neuroimmune response to tick bites that leads to a scratching response and consequently tick removal. In contrast, mice, including natural reservoirs or strains used in preclinical models, generate a skin immune response to tick bites but without active tick removal (*46*). Then it appears that tick-sensitized mice do not scratch in response to tick bites and do not mechanically remove ticks even in the presence of a skin immune response (*46*). This indicates that the active neurological component that triggers IITR and the scratching response to tick bites may be absent in mice due to their distinct neuroimmune circuits. The mouse model is therefore ideal for the study of ATR, characterized by a basophilic response and anti-tick antibodies that directly affect tick fitness (*47*). Alternatively, we propose GPs as a model to study itch responses to blood feeding arthropods, providing an opportunity to investigate the mechanisms responsible for ectoparasite sensing and removal.

A short time, 5 days from initial tick exposure, is required to develop the neuroimmune response that leads to IITR. This suggests that an intervention based on itch-induction may be effective in preventing pathogen transmission shortly after immunization. This is in contrast to ATR which has been shown to require between 60 to 90 days after initial tick exposure for its development (*14*). Importantly, our demonstration that a single tick can prime the development of this fast neuroimmune response to tick bites suggests that a small dose of itch-inducing antigen/s will be sufficient for an effective anti-tick vaccine. The rapid subsiding of itch post tick removal is another advantage for a vaccine as it precludes unnecessary discomfort, or even damage to the host skin.

Our study provides evidence that itch serves as a defensive mechanism for the removal of ectoparasites, proposing a potential basis for the evolutionary conservation of itch. It will be important to determine whether similar adaptive responses are employed for defense against other ectoparasites. Although we hypothesize that ectoparasite removal is the major evolutionary drive for the development of itch, our results do not exclude contribution by other potential mediators such as inflammation (*48*). Similarly, our results do not exclude other non-itch mediated processes being involved in defense against ectoparasites.

In summary, we provide evidence for the presence of IITR. Driven by an early neuroimmune response to tick bites in a tick-sensitized host, IITR causes localized and temporary itching and a fast mechanical tick removal and occurs in a window of time that precedes transmission of many tick-borne pathogens. Further understanding of this neuroimmune response may lead to the identification of itch-inducing tick antigens that can be developed as an innovative “behavioral assisted” anti-tick vaccine.

## MATERIAL AND METHODS

### Ethics statement

All experimental animal procedures were reviewed and approved by the National Institute of Allergy and Infectious Diseases (NIAID) Animal Care and Use Committee (ACUC) under animal protocol ASP LMVR6. The NIAID DIR Animal Care and Use Program complies with the Guide for the Care and Use of Laboratory Animals and with the NIH Office of Animal Care and Use and Animal Research Advisory Committee guidelines. Detailed NIH Animal Research Guidelines can be accessed at https://oma1.od.nih.gov/manualchapters/intramural/3040-2/.

Efforts were made in the planning and execution of all animal procedures to comply with the guidelines of the National Center for Replacement, Refinement and Reduction (3Rs) of Animals in Research (https://nc3rs.org.uk/the-3rs).

Human skin samples were obtained from participants enrolled onto protocol NCT05036707. The study is approved by the institutional review board (IRB) at the National Institutes of Health (NIH) (Bethesda, MD). All participants signed written informed consent.

### Guinea pigs

3-6 weeks old out-bred female or male albino Hartley guinea pigs, *Cavia porcellus* (Stain code: 051, https://www.criver.com/products-services/find-model/hartley-guinea-pig?region=3611), were obtained from Charles Rivers laboratories, Wilmington, MA, USA and were communally housed in groups of up to three animals under static cage conditions at the NIAID Twinbrook animal facility, Rockville, MD.

### Ticks

Pathogen-free *Ixodes scapularis* nymphal ticks were either obtained from the Centralized Tick Rearing Facility, Department of Entomology and Plant Pathology, Oklahoma State University, Stillwater, OK, USA, (main source) or sourced from the established *Ixodes scapularis* colonies at the Rocky Mountain Laboratories (RML), NIAID, Hamilton, MT, USA (secondary source) and the Laboratory for Malaria and Vector Research (LMVR), NIAID, NIH, Rockville, MD, USA (temporary source). Nymphal ticks were housed in cotton-wool-sealed plastic tubes or gauze-mesh-sealed universal tubes both sealed in zip-lock bags containing a moist sponge for humidity. These bags were kept for long-term storage in a housing incubator at 20°C and ≥95% humidity under a 16:8 h photoperiod at the Laboratory for Malaria and Vector Research (LMVR), NIAID, NIH, Rockville, MD, USA.

### Experimental guinea pig exposures to nymphal ticks

For experimental use, nymphal ticks were transferred from the cool housing incubator to a warm (∼27°C / 95-98% humidity) incubator up to 4 days prior to experimental use. To break diapause and for acclimatization, zip-lock bags containing nymphal ticks were transferred to a windowsill for sunlight exposure at room temperature 1-2 days prior to experimental use and left there until use.

For nymphal tick exposures, the head and neck area of GPs were shaven (not too closely) with electrical clippers, leaving 1-2 mm stubbles. Our nymphal ticks preferred the stubble over depilated skin as it facilitated holding on to the host and stimulated their sensilla once they burrowed in between the stubble. GPs were then anesthetized individually in 5 – 7 min intervals by intraperitoneal (i.p.) administration of a Ketamine (50 mg/kg) / Xylazine (5 mg/kg) mixture according to body weight. Using a fine paint brush, nymphal ticks were stimulated to burrow in between the remaining hair stubbles by repeatedly tapping them lightly in intervals during the first 10 – 15 min post placement. By brushing hard against the grain of the hair stubble, nymphal ticks that would not attach to the host skin were identified and replaced with new nymphal ticks. This cycle was repeated until the desired number of ticks had attached. This procedure increased the likelihood of tick attachment at a target location to >95%. GPs were placed for recovery (60 – 90 min post induction) in triple containment cages, where the outermost cage contained a water barrier and rims coated with Vaseline to prevent tick escape. ∼3 h post tick attachment (pta), GPs were anesthetized with isoflurane (Forane by Baxter, Deerfield, IL, USA) gas (4% at 2 L/min O_2_ flowrate), and attached nymphal ticks were counted. If applicable, attached nymphal tick numbers were reduced to a desired number for all GPs in an experiment. GPs were also checked for nymphal ticks that had bitten off-target, which were removed, too. In some experimental tick exposures, Elizabethan collars (e-collars; Lomir Biomedical Inc., Malone, NY, USA) were placed around the GPs’ neck to prevent premature tick removal by the GPs and approximately equal antigen delivery. Tick-exposed animals were also housed individually to prevent social grooming. At the end of experimental tick exposures, remaining nymphal ticks were removed with tweezers and GPs were returned to communal housing cages.

The maximum permitted duration of nymphal tick attachment during a nymphal tick exposure varied between experiments. In general, GPs were sensitized by weekly nymphal tick exposures lasting 24 to 72 hours each as required and resting periods of 6 and 4 days, respectively. Only in one experiment, 18 days of rest were observed between exposures. Depending on the experiment, remaining nymphal ticks were counted at 12 hours pta, 24 hours pta, 36 hours, 48 hours pta, and/or 72 hours pta

### Tick health assessment

Tick-naïve and three (3) times sensitized GPs were exposed to ticks as described above and e-collars were applied at 3 hours pta after tick adjustments to 15 attached nymphal ticks. At 48 hours and 72 hours pta attached nymphal ticks were recovered from half of the exposed GPs in each group, respectively. Care was taken to remove nymphal ticks by pulling gently with fine forceps to leave the hypostome intact. Recovered nymphal ticks were immobilized on double sided tape and imaged under a stereomicroscope (Leica M165 FC) with a Leica DFC 7000 T camera (Leica, IL, USA) using the Leica Application Suite X v.3.7.4.23463, which was also used for measurements of the length of the nymphal alloscutum’s distention and the scutum’s width (fig. S1D). These measurements were used to calculate the scutal index, which is a ratio produced by dividing the alloscutum’s length by the scutum’s width (*49*). The scutal index is an established indicator of tick feeding success, in particular, in instances when ticks are not allowed to feed to repletion or non-adult tick stages (here, nymphs) are used, excluding assessment of tick molting or egg mass. The concept is that length measurement of the alloscutum will increase with continued feeding over time, while the scutum remains unchanged during the feeding and thus, its width is fixed. Thus, the larger the ratio, the greater the engorgement. Further, nymphal ticks were visually assessed for intactness and movement under the stereomicroscope as indicators of nymphal tick life-status; visible intactness and movement = alive, no movement despite stimulation by tapping with a fine paint brush or visible damage to/desiccation of the tick = dead.

### Scratch bout quantification

We employed PhenoTyper^®^ home cages with video tracking technology using EthoVision^®^ XT v.17 for video analysis (Noldus Information Technology INC, VA, USA) to assess changes in guinea pig behavior post tick attachment. We employed an infrared camera system to record videos overnight in total darkness. GPs are often described as crepuscular, but studies have shown that their sleep occurs in short bursts and is evenly distributed over 24 hours (*50*) As animals were recorded for 14 h, we provided custom-made absorbent cage liners (Top Layer: 85 gsm brushed/knitted polyester; Middle Layer: 250gsm Polyester Soaker; Bottom Layer: 85 gsm brushed/knitted polyester bottom with waterproof TPU finished top) that were dyed with an infrared absorbent dye, which gave the liners a black appearance under the infrared camera. Experimentally, we exposed GPs to nymphal ticks as described above and either recovered them from the ketamine/xylazine general anesthesia in the PhentoTyper^®^ home cages after tick placement or moved them into the PhentoTyper^®^ home cages ∼5 h after nymphal tick placement. Black curtains were drawn around the cage to reduce stray room light and obstruct guinea pig vision of the room to avoid behavioral freezing. In EthoVisionXT, video tracking was conducted post recording and commenced when, firstly, the center point of a guinea pig was detected for ≥5 sec, and, secondly, the GPs’ activity exceeded 0.05%, which indicated presence in the cage or recovery from anesthesia. Video tracking covered the first 12 hours post-anesthesia recovery or presence in the cage depending on experiment. Scratch bouts to the top of the head were used as an indicator of nymphal tick-induced itch, according to the definition established by Shimada & LaMotte (2008) (*51*). For the video analysis we defined potential scratch bouts in EthoVision^®^ XT by combining two “multi condition” criterions, consisting of the following criteria: a) “body elongation” (threshold: <47%, Average: 10), “activity” (≥0.1%, Average: 10), and b) “body elongation” (threshold: <46.5%, Average: 10), “distance moved” (nose-point: <0.58 cm; center-point: <0.3 cm; Track Smoothing (Lowess): 9), “activity” (≥0.26%, Average: 10), “velocity” (nose-point: <12 cm/s; center-point: <5 cm/s; Average: 10), “in zone” (Body points: nose point; not in the following zone: feeder zone). The output data was screened in R statistical software to narrow possible scratching events by excluding all events shorter than a) 1 sec, and b) 0.66 sec, respectively. Further, all events that showed an overlap of <20% of a) and b) were excluded. All filter events were manually verified and scored in EthoVision^®^ XT and used as the basis to quantify scratch bouts (duration and frequency) for the statistical analysis.

### Enzyme-Linked Immunosorbent Assay (ELISA)

Detection of anti-*Ixodes scapularis* IgG and IgE in tick-naïve and weekly exposed GPs for three (animals from week 1, week 2, week 3 and week 4 groups) or four times (animals from week 10 group), were analyzed by indirect ELISA IgG using *I. scapularis* salivary gland homogenate (SGH), and a commercial IgE kit (MyBioSource, Inc., San Diego, CA, USA). Briefly, for IgG assay, microtiter plates were coated with 5 ug/mL of SGH and incubated overnight at 4 °C. Subsequently, plates were blocked with PBS/10% FBS during 1 hour at room temperature. Plates were washed six times with PBS/0.05% Tween-20 and incubated with serum diluted 1:500 in PBS/5% FBS in duplicate. After 1 h at room temperature, plates were washed six times, and wells were filled with the secondary detection antibody (Goat anti-Guinea Pig IgG (H+L) – Invitrogen - diluted to 1:5,000 in PBS/ 5% FBS). The plates were incubated for 1 h at room temperature. After twelve washes, the 1-Step ABTS substrate solution was added. The optical density (OD) was measured at 405 nm, using a 96-well microplate reader after 30 min. ELISA IgE assay was performed following the manufacturer’s recommendation. Briefly, guinea pig sera were diluted 1:16 and spiked-in samples were used as control (final concentration 50 ng/mL). After 15 minutes, HRP reaction was stopped, and the OD was measured at 450 nm. The IgE concentration was assessed using a standard curve and the interpolated values were obtained using a linear regression calculated with GraphPad Prism software.

### Disruption of establishment of acquired tick removal

Fingolimod (FTY720) is an immunomodulator drug that modulates the sphingosine 1-phosphate (S1P) receptors on naïve lymphocytes preventing their egress from lymph nodes (*35, 52*). Five mg of FTY720 (Sigma-Aldrich) were suspended in 3 ml of PBS (Lonza. Portsmouth, NH) under sterile conditions by pipetting the suspension up and down several times. The suspension was transferred to a 15 ml Falcon tube and cycled 3 times through 10 min heating in a water bath at 37°C followed by vortexing for 10 min at 2,000 rpm at room temperature. Tick-naïve GPs were treated with 4 mg/kg of the FTY720 suspension starting 5 – 7 days prior to the first exposure. Circulating lymphocytes were counted pre-treatment and the day before exposure to assess FTY720-induced reduction in circulating lymphocytes. Treatment was continued until the end of the experiment. Nymphal tick exposures were conducted as described above.

### Resiniferatoxin administration

Resiniferatoxin (RTX) (1mg) was obtained from AdipoGen Life Sciences (CA, USA). RTX was solubilized in 200 μL 100% ethanol to make a RTX stock concentration of 5μg/μL. It was further diluted to a working concentration of 100ng/μL RTX in sterile vehicle (0.25% Tween 80, 2mM ascorbic acid, 0.9% NaCl). Intrathecal injections (2mg RTX in 20μL working solution) in guinea pig were made using a 500 μL zero dead volume insulin syringe with a fixed needle of 29G (MHC Medical Products, OH, USA). To test for loss of Trpv1-neuron activity, intraplantar administration of 20 ug Capsaicin (Cat#M2028, Millipore Sigma, USA) in 20 μL of vehicle solution (5% Tween 80, 5% Ethanol, 0.9% NaCl) was made 7 days after RTX injection and number of licking bouts counted.

### Transcriptome sequencing and analysis

Total RNA was extracted from skin biopsies as previously described (*53*). Briefly, biopsies were stored in DNA/RNA Shield (Zymo research) and kept at 4 °C until processing. Skin was homogenized using 3-mm zirconium beads (Biolink laboratories) and a Magnalyser bead beater (Roche Diagnostics®). After homogenization total RNA was loaded at InnuPure® C16 touch (AnalitikJena) for magnetic separation of total RNA using an InnuPREP® RNA Kit–IPC16 (AnalitikJena). Total RNA was sent for library preparation and sequencing (Novogene Co.). mRNA was enriched by Poly (A) capture and RNA libraries were prepared using the NEBNext Ultra II Directional RNA Library Prep Kit for Illumina (New England Biolabs) and sequenced on a NovaSeq 6000 (Illumina) generating paired end reads at 150 bp length. Raw RNA sequences were prepared and analyzed using the CLC genomics workbench v22 (Qiagen). Sequences were trimmed (prepare raw data workflow) to remove poor quality sequences and adaptors. Trimmed reads were mapped and the differential expression of genes for the groups (3 times sensitized Tick vs one-time sensitized vs unbitten skin) was calculated for 4 biologically independent samples (RNA-seq and differential gene expression analysis workflow). NCBI Cavpor3.0 assembly and annotation were used as a reference genome. As previously described (*54*), filtering was performed on all mapped gene counts to exclude genes where the sum of counts in all conditions was inferior to 10. Significant associations were considered when a p <0.01 and log2 fold change larger than 1.5 (+/-). The principal component analysis (PCA) was performed with log2 TPMs +1. Heatmap was generated with log2 TPMs +1 z-scores. Heatmap and PCA plots were generated using the GraphBio web app68.

### Laser Speckle

Laser speckle contrast imaging is a non-invasive technology that detects the dynamic change in backscattered light (cellular motion increases light scatter) (*55*). It is used in clinical settings to assess perfusion of various tissues, which includes cellular influx into sites of skin inflammation. GPs were tick exposed as described above. 24 hours pta, GPs were anesthetized by isoflurane delivered by a SomnoFlo vaporizer (Kent Scientific, Torrington, CT, USA) and hair around tick-bite sites (top of the head) was removed by Nair (Church & Dwight Co., Inc., Ewing, NJ, USA) treatment. Briefly, Nair was applied according to supplier’s instructions for 3 to 5 min (longer treatment times were detrimental to tick survival). Nair was washed off under running water. The following day, GPs were anesthetized individually by i.p. administration of a Ketamine (50 mg/kg) / Xylazine (5 mg/kg) mixture according to body weight and placed on a PhysioSuite heating pad (Kent Scientific, Torrington, CT, USA). Nymphal ticks attached to anesthetized GPs were visualized with a PerCam PSI HR (Perimed, NV, USA) calibrated using a LI723 CalBox PSI HR (Perimed) and the PIMsoft package v1.5 (Perimed). 1 cm^2^ skin areas were recorded for 1 min at a time. Circular regions of interest (ROI; 3.47 mm^2^) were drawn around each tick (excluding the tick itself, as it masked the signal) and one unbitten area per image for background signal measurements. Measurements were recorded for 1 min and subsequently averaged (smoothed) over a time of interest (TOI) of a 20 s range of the 1 min recoding. All readings were normalized by subtracting the background value from the measurements around the tick by the respective image, producing Δ perfusion units.

### Histology

Skin biopsies of nymphal tick bitten, and unbitten skin (ø 3 mm) were collected by punch biopsy from CO_2_ euthanized GPs. Fresh biopsies were immediately placed in 10% neutral formalin and incubated 24 hours at 4°C for thorough tissue fixation. Fixed skin biopsies were washed in PBS, stored in 70% ethanol, and submitted to Histoserv, Inc. (Histoserv, Inc., MD, USA) for paraffin embedding, sectioning, and staining with hematoxylin and eosin (H&E; provided by Histoserv). All paraffin skin sections were cut at 5 µm thickness. Samples were collected from tick-naïve, immunosuppressed (FTY720) and tick-sensitized GPs. Stained slides were digitalized using a Motic Easyscan Pro 24 slide scanner. H&E images where processed and quantified for cellular infiltrate using Qupath software v0.5.1 (*56*). Cell detection workflow was applied to IHC images. Machine learning object classification by random trees was applied on skin biopsies scans with at least 20 points/category. Cell detection workflow was applied to all images after algorithm training.

### Immunohistochemistry

Formalin-fixed paraffin-embedded (FFPE) tissues sections (5 µm thickness) obtained from guinea pig skin were used to perform immunohistochemical (IHC) staining using the following antibodies: a rat monoclonal CD3 with colon CD3-12 (AbD Serotec, Catalog No: MCA1477, dilution of 1:600) and a rabbit polyclonal Iba1 (Wako, Catalog No: 019-19741, dilution of 1:800). Staining was carried out on the Bond RX (Leica Biosystems) platform according to manufacturer-supplied protocols. Briefly, 5 µm sections were deparaffinized and rehydrated. Heat-induced epitope retrieval (HIER) was performed using Epitope Retrieval Solution 1, pH 6.0, heated to 100°C for 20 minutes. The specimen was then incubated with hydrogen peroxide to quench endogenous peroxidase activity prior to applying the primary antibody. Detection with DAB chromogen was completed using the Bond Polymer Refine Detection kit (Leica Biosystems, Catalog # DS9800). Slides were finally cleared through gradient alcohol and xylene washes prior to mounting and placing cover slips. Sections were examined by a board-certified veterinary pathologist (DAA). Immunohistochemistry images where processed and quantified using Qupath software v0.5.1 (*56*). For macrophage staining (Iba1), ‘Cytoplasm: DAB OD mean’ was the score compartment, with a threshold of .05 for intensity parameters. For T-cell staining (CD3), ‘Nucleus: DAB OD mean’ was the detection setting and the threshold was 0.1 for intensity parameters. All other settings were left as default. ROIs were determined based on the area of highest infiltration for the largest biopsy analyzed for each figure.

### In situ hybridization and imaging

Fresh frozen GP DRGs were sectioned at 20 µm with a cryostat. Multiplex in situ hybridization was performed with RNAscope® multiplex fluorescence assay (Advanced Cell Diagnostics, Inc). Guinea pig spinal cords or dorsal root ganglia were embedded into cryostat embedding media, freshly frozen, and then cryosectioned at 20-μm thickness. RNAscope probes for Nppa (REF#1555751-C1), Tubb3 (REF#1556171-C2), Trpv1 (REF#1231081-C3), Npr1 (REF#1574901-C1), Il31ra (REF#1574911-C2) and Osmr (REF#1231091-C1) were included in this study. Fluorescence images were collected with Nikon Eclipse Ti confocal laser-scanning microscope.

### Exposure of human volunteers to *I. scapularis* ticks

National Institutes of Health (NIH) protocol NCT05036707 is an ongoing study exposing human research volunteers to laboratory-reared, pathogen-free, larval *I. scapularis* ticks. The protocol is under an investigational device exemption (IDE) approved by the Food and Drug Administration, conducted in accordance with Good Clinical Practice guidelines, and a Sponsor medical monitor reviews interim safety data. All participants signed NIH institutional review board approved informed consent, had no clinical history of tickborne diseases and no known history of tick bites. A small cohort of 4 research volunteers had samples available 24 hour or 48 hours after first, second or third tick exposure and were included in this analysis. Control skin biopsy was collected on Day 0 prior to tick exposure. Ten laboratory-reared larval ticks were placed at 2 different body sites (total of 20 ticks). Tick colony description and tick exposure procedure are described in Marques et al. (*57*). Research volunteers returned in 24 hours to remove ticks from one of the body sites and to collect skin biopsy after the first tick placement. Research volunteers returned at 24 or 48 hours after second and third placements to remove ticks from the second body site and to collect skin biopsy.

### Statistical analysis and figure panel generation

All statistics were conducted in RStudio v.2024.04.1-748 (*58*) using R v.4.4.3 (*59*) with the exception of bulk RNAseq analysis and the neurological itch marker expression data. All data graphs, presented in the main figures and extended data figures were mainly generated in GraphPad PRISM v. 10. 2.3 (*60*) and occasionally in Adobe Illustrator v. 29.4 (64-bit; https://www.adobe.com/products/illustrator.html). Experimental schematics were generated in BioRender (Fig. 1B: https://BioRender.com/2jxmm9k; fig. S1A: https://BioRender.com/47zaovj; fig. S5A: https://BioRender.com/rduvqd2; fig. S6A: https://BioRender.com/dtivqaj; fig. S8A: https://BioRender.com/y2smy29). For details on the applied statistics, please, consult the detailed statistics report in the supplementary materials to this publication. All R scripts underlying the statistical and data analysis are available at https://github.com/joedoehl/Immune-Induced-Itch-Drives-Rapid-Tick-Removal-in-Vertebrates.git.

## Acknowledgements

We would like to thank Yvonne Rangel Gonzalez and Brittany Mills from VMBS, NIAID for technical support. We would like to thank the Twinbrook 3 animal facility personnel for their support and professionalism.

## Note

This article reflects the views of the author and should not be construed to represent FDA’s views or policies. The content of this publication does not necessarily reflect the views or policies of the Department of Health and Human Services, nor does mention of trade names, commercial products, or organizations imply endorsement by the U.S. Government.

This research was supported by the Intramural Research Program of the National Institutes of Health (NIH). The contributions of the NIH author(s) were made as part of their official duties as NIH federal employees, are in compliance with agency policy requirements, and are considered Works of the United States Government. However, the findings and conclusions presented in this paper are those of the author(s) and do not necessarily reflect the views of the NIH or the U.S. Department of Health and Human Services.

## Author contributions

Conceptualization: J.G.V., J.M.C.R., M.A.H., J.S.P.D., T.D.S., L.T., A.M. Funding acquisition: J.G.V., M.A.H., J.M.C.R., D.A.A., L.T., A.M.; Investigation: J.S.P.D., T.D.S., S.D., C.S.G., E.I., L.R., R.D., P.C., R.F., X.G., P.T., A.D.S.M., M.M., J.O., H.A., S.B., C.P., S.T., D.A.A., J.M.A., D.E.S., J.G.V. : Methodology: J.S.P.D. T.D.S., S.D., C.S.G., E.I., L.R., R.D., P.C., R.F., X.G., P.T., A.D.S.M., M.M., J.O., H.A., S.B., C.P., S.T., D.A.A., J.M.A., D.E.S.; Project administration: J.G.V. Supervision: J.G.V., M.A.H., J.S.P.D., T.D.S., D.A.A., A.M.; Visualization: J.S.P.D., T.D.S, D.A.A., J.G.V., R.F., L.R., M.M. Writing – original draft: J.G.V., J.S.P.D., D.E.S., T.D.S., M.A.H. Writing – review & editing: J.G.V. J.M.C.R., M.A.H., S.K., F.O. J.S.P.D., T.D.S., L.T., A.M., S.D., C.S.G., E.I., L.R., R.D., P.C., R.F., X.G., P.T., A.D.S.M., M.M., J.O., H.A., S.B., C.P., S.T., D.A.A., J.M.A., D.E.S., J.G.V.

## Material availability

This study does not generate new unique reagents. All data are available in the manuscript or the supplementary materials. Materials described in this manuscript are available from J.G.V. on request.

## Data and code availability

All raw and processed sequencing data are available at NCBI, BioProject ID: PRJNA1259463, Reviewer link: https://dataview.ncbi.nlm.nih.gov/object/PRJNA1259463?reviewer=r47cm3ftc888ia8giu1s072o <u>be</u>. All data underlying the statistical analyses in the supplementary statistics report is also available from the author’s GitHub repository: https://github.com/joedoehl/Immune-Induced-Itch-Drives-Rapid-Tick-Removal-in-Vertebrates.git.

---

**Figure S1.**
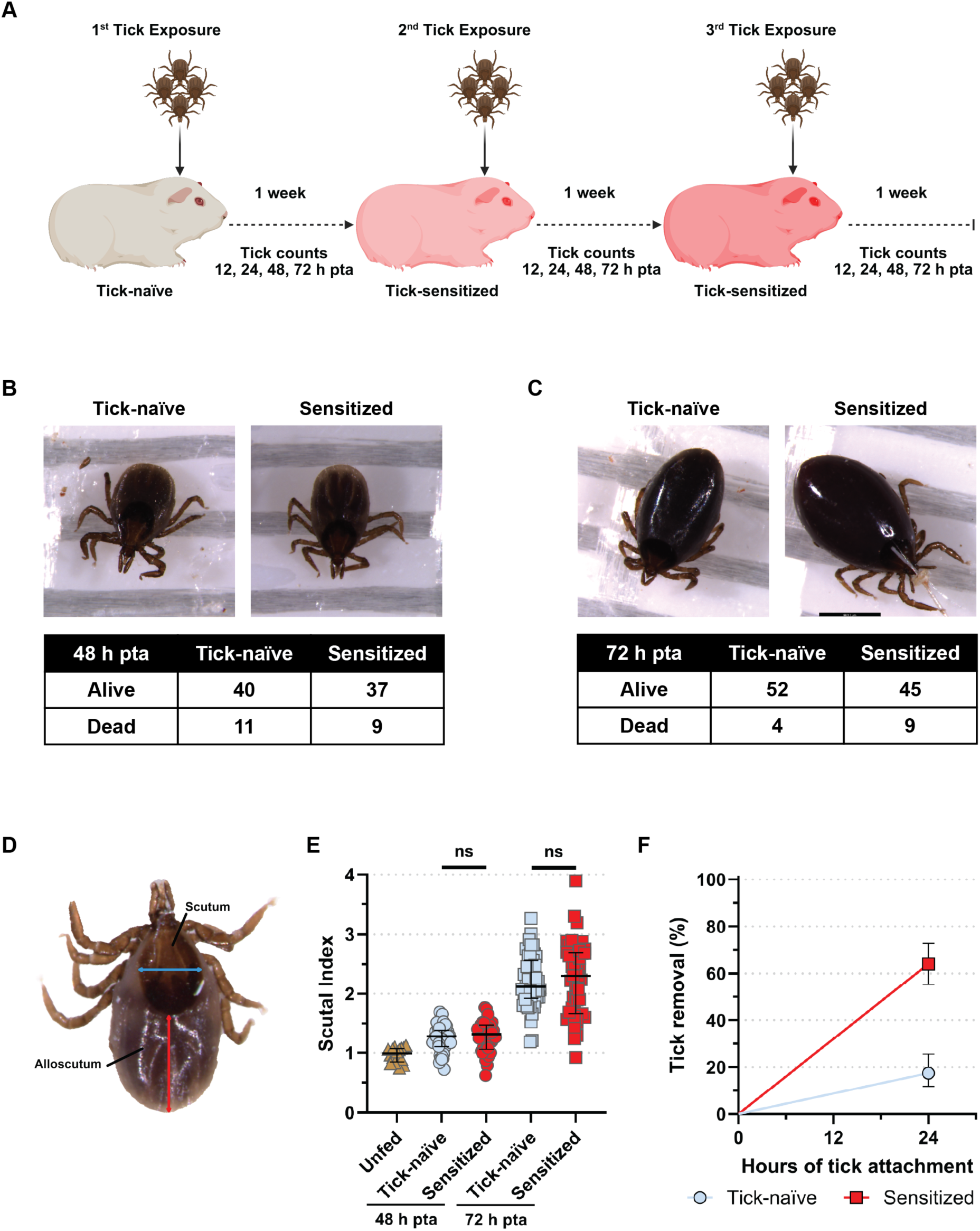
(**A**) Repeated tick exposure experimental design (Created in BioRender: https://BioRender.com/47zaovj). (**B,C**) Images of ticks fed on sensitized and tick-naïve guinea pigs (GPs) with a respective cross table of live/dead tick counts at 48 and 72 h post tick attachment (pta). Cross table analysis by (**B**) Chi^2^ test (p=1), and (**C**) Fisher’s Exact test (p=0.5124). (**D**) The scutal index is a ratio produced by dividing the alloscutum’s length by the scutum’s width. (**E**) Scatter plot: Scutal index values of nymphal ticks fed on tick-naïve (48 h pta: *N*=42, 72 h pta: *N*=50) and sensitized (48 h pta: *N*=38, 72 h pta: *N*=48) GPs, or unfed (*N*=16). Two-way ANOVA (excluding unfed): Sensitization state (tick-naïve / tick sensitized) p=0.3229, Time (48 h pta versus 72 h pta) p<0.0001. Median ± IQR shown. (**F**) Smoothed, inverted Kaplan-Meyer plot showing probability of tick removal (%) during a 1^st^ (Tick-naïve, *N*=120 ticks) and a 4^th^ (Sensitized, *N*=114 ticks) tick-exposure at 24 h pta. Analysis by Peto & Peto modification of the Gehan-Wilcoxon test: p<0.0001. 95% CI shown. For detailed statistics see supplementary report.

**Figure S2.**
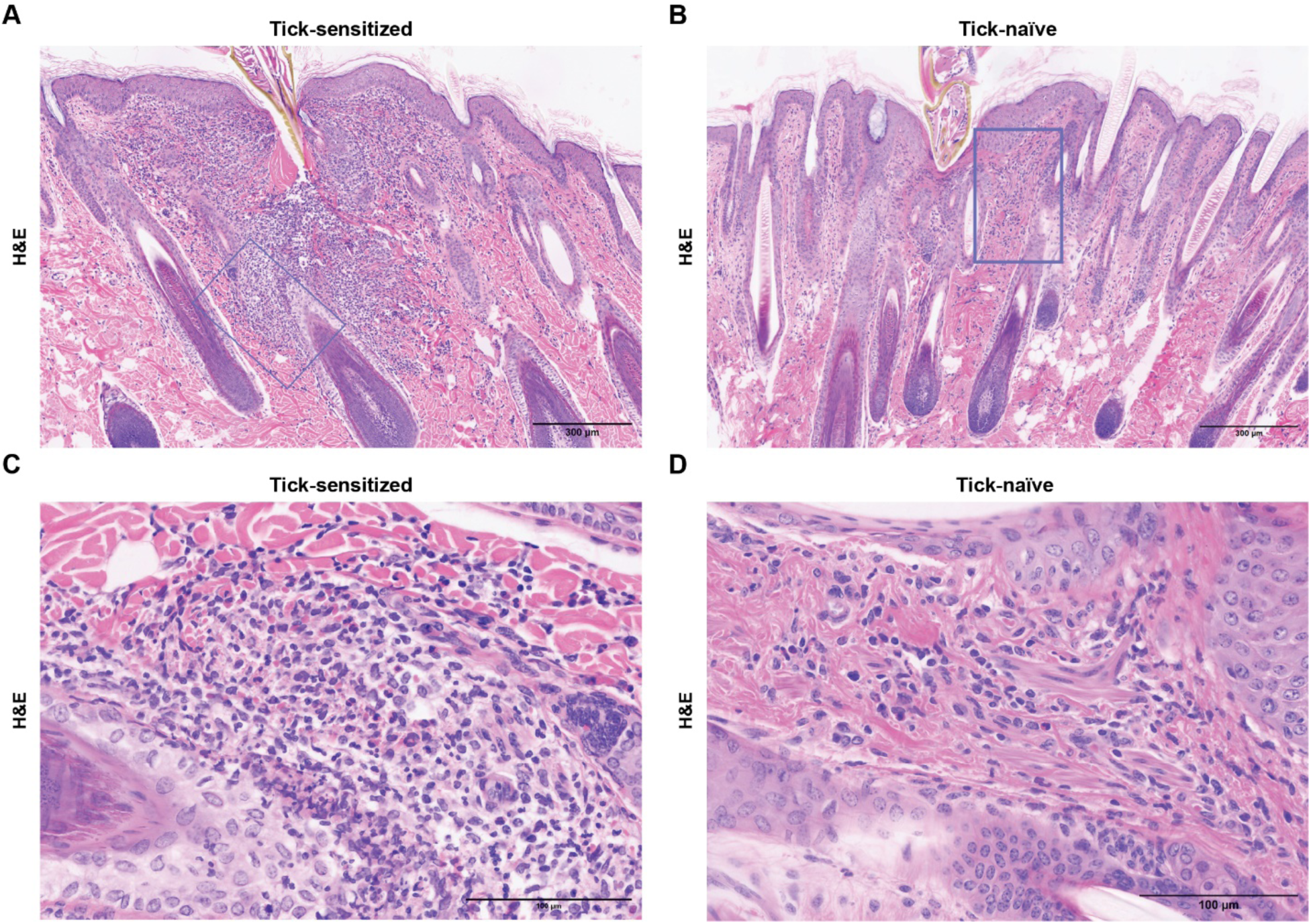
(**A,B**) Representative H&E skin cross-sections (>|5µm|<) of tick bite sites 24 h post tick attachment (pta) in (**A**) three times weekly tick-sensitized and (**B**) first-time exposed tick- naïve guinea pigs (GPs). (**A**) Significant focally extensive dermal inflammation consisting primarily of a mononuclear cell infiltrate surrounding and separating adnexal structures. (**B**) Mild epidermal hyperplasia with a minimal dermal infiltrate. (**A,B**) Scale bar: 300 µm. 10x (**C,D**) Enlarged section from (**A,B**) to illustrated (**A**) presence and (**B**) absence of eosinophils in tick bite sites. Scale bar: 100 µm.

**Figure S3.**
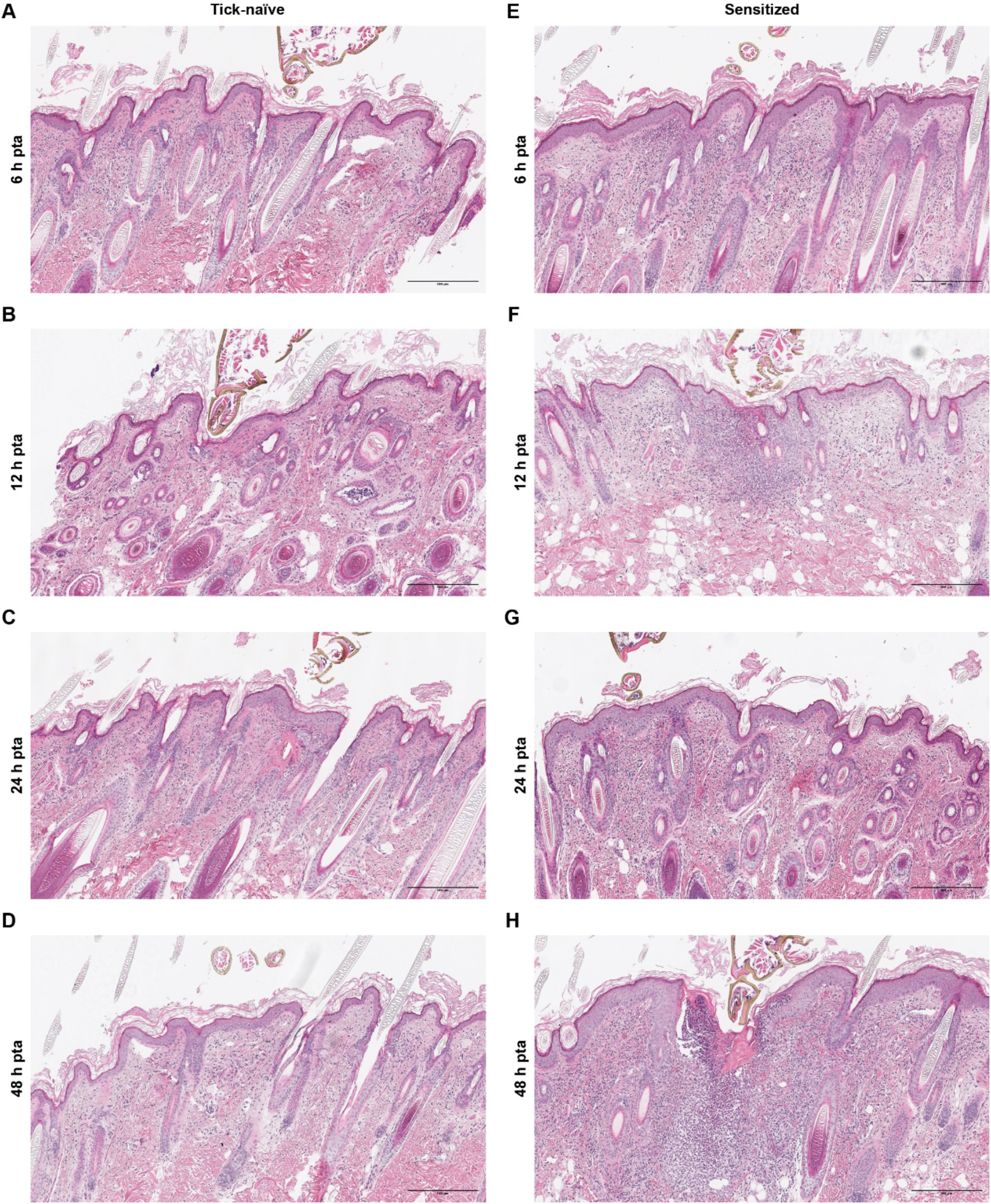
Representative histological sections (>|5µm|<) stained with H&E showing cell infiltrates at the tick bite sites in tick-naïve and tick-sensitized GPs over 48 h pta. Scale bar 300 µm. 10x.

**Figure S4.**
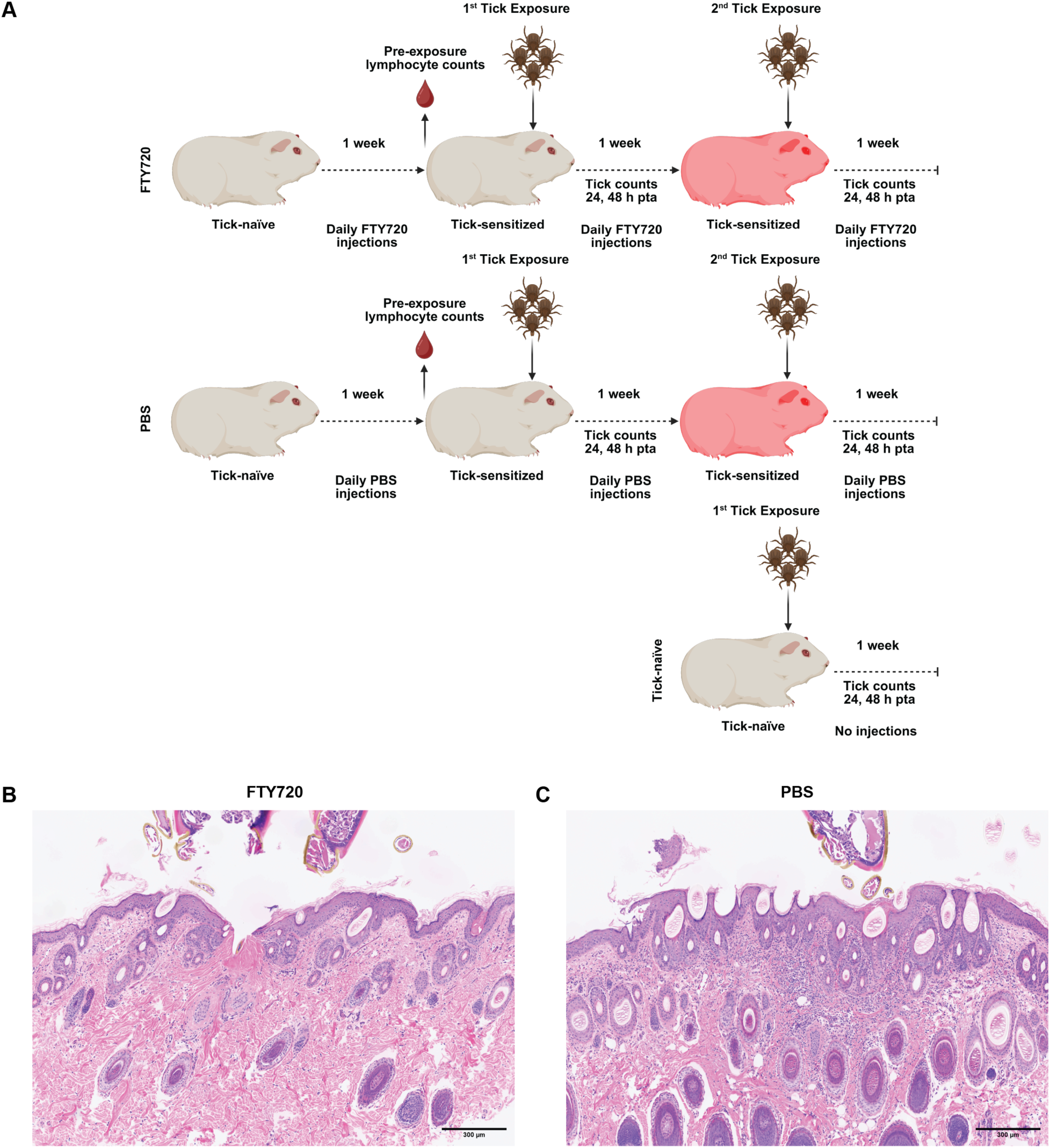
(**A**) FTY720 experimental design (Created in BioRender: https://BioRender.com/rduvqd2). Guinea pigs (GPs, *N*=10/group) were either FTY720- or PBS-treated and then exposed two times to 15 ticks each. First-time exposed tick-naïve GPs were used as a tick attachment control. (**B, C**) Representative H&E skin cross-sections (>|5µm|<) of a dermal biopsy at tick bite sites 48 h pta during the second tick exposure. 10x. (**B**) FTY720- treated GP showing mild epidermal hyperplasia and dermal edema but a minimal dermal infiltrate reminiscent of a first-time infested tick-naïve GP. (**C**) PBS-treated GP showing significant dermal inflammation and epidermal hyperplasia.

**Figure S5.**
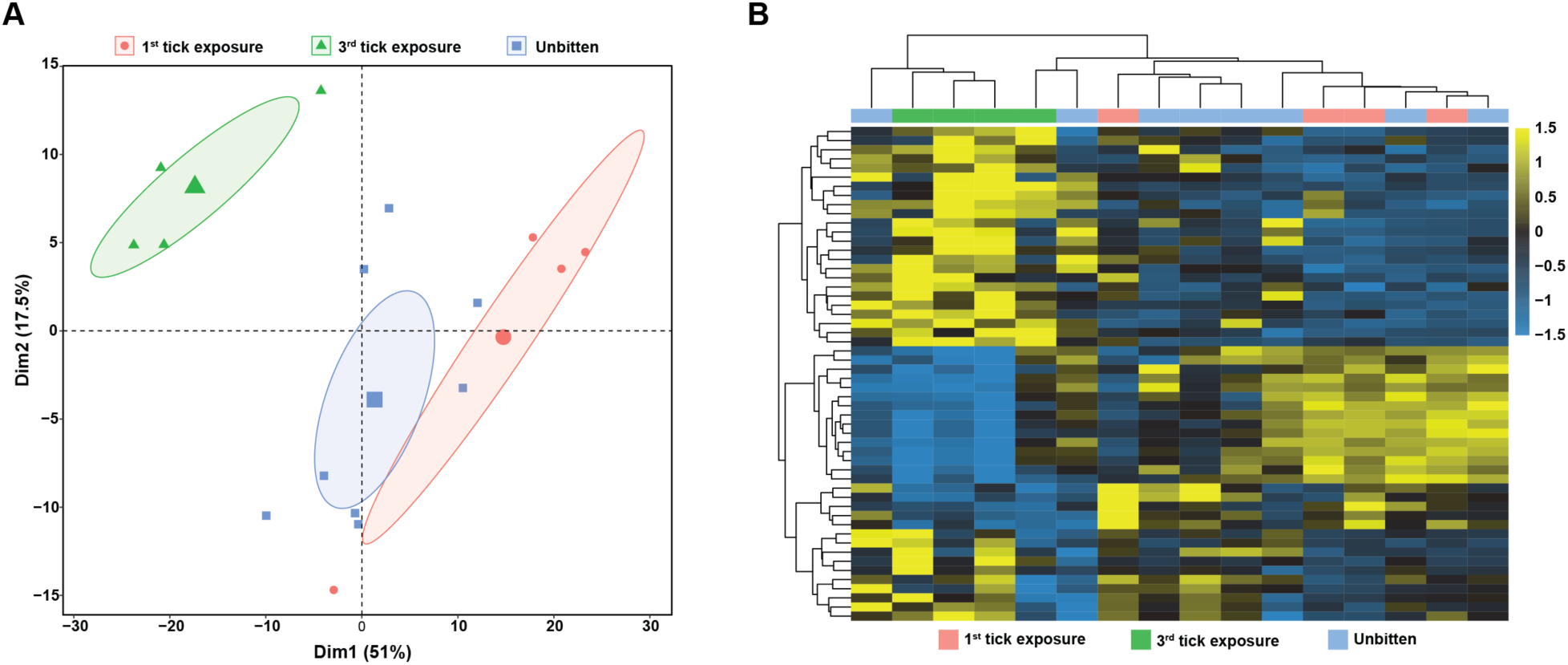
RNAseq analysis of 3 mm skin biopsies from tick bite sites during a first (*N*=4) and third (*N*=4) tick exposure, and unbitten skin (*N*=8). (**A**) PCA plot. (**B**) heatmap of the top 54 differentially expressed genes, either upregulated or downregulated, after applying a ±3 log2 fold change cutoff.

**Figure S6.**
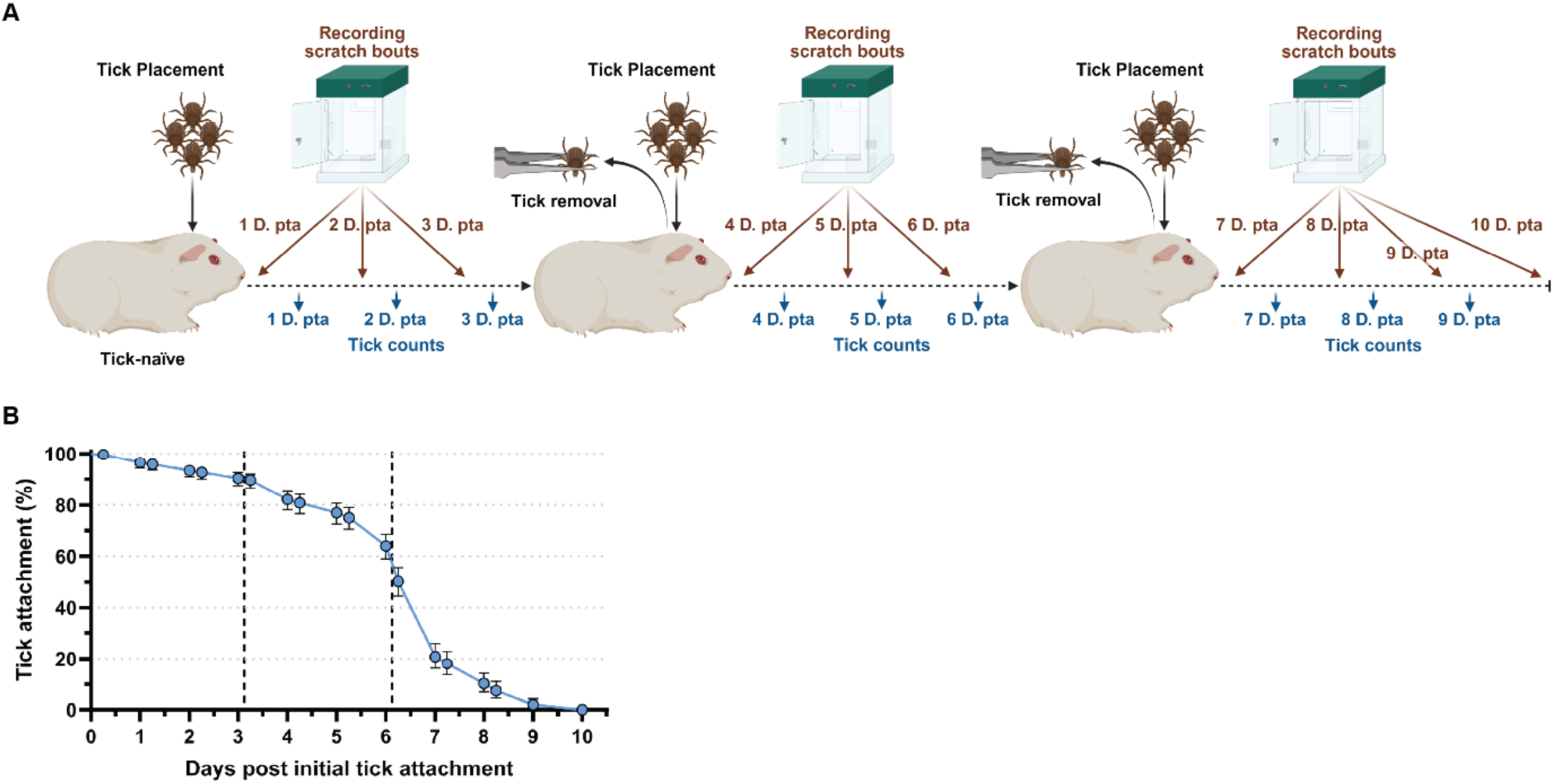
(**A**) Experimental design schematic of continuous tick exposure (Created in BioRender: https://BioRender.com/dtivqaj). (**B**) Smoothed Kaplan-Meyer plot showing probability of tick attachment (%) to continuously tick-exposed tick-naïve GPs up to 10 days post initial tick attachment (*N*=10 GPs). Dashed vertical lines indicate re-placement of ticks. 95% CI shown.

**Figure S7.**
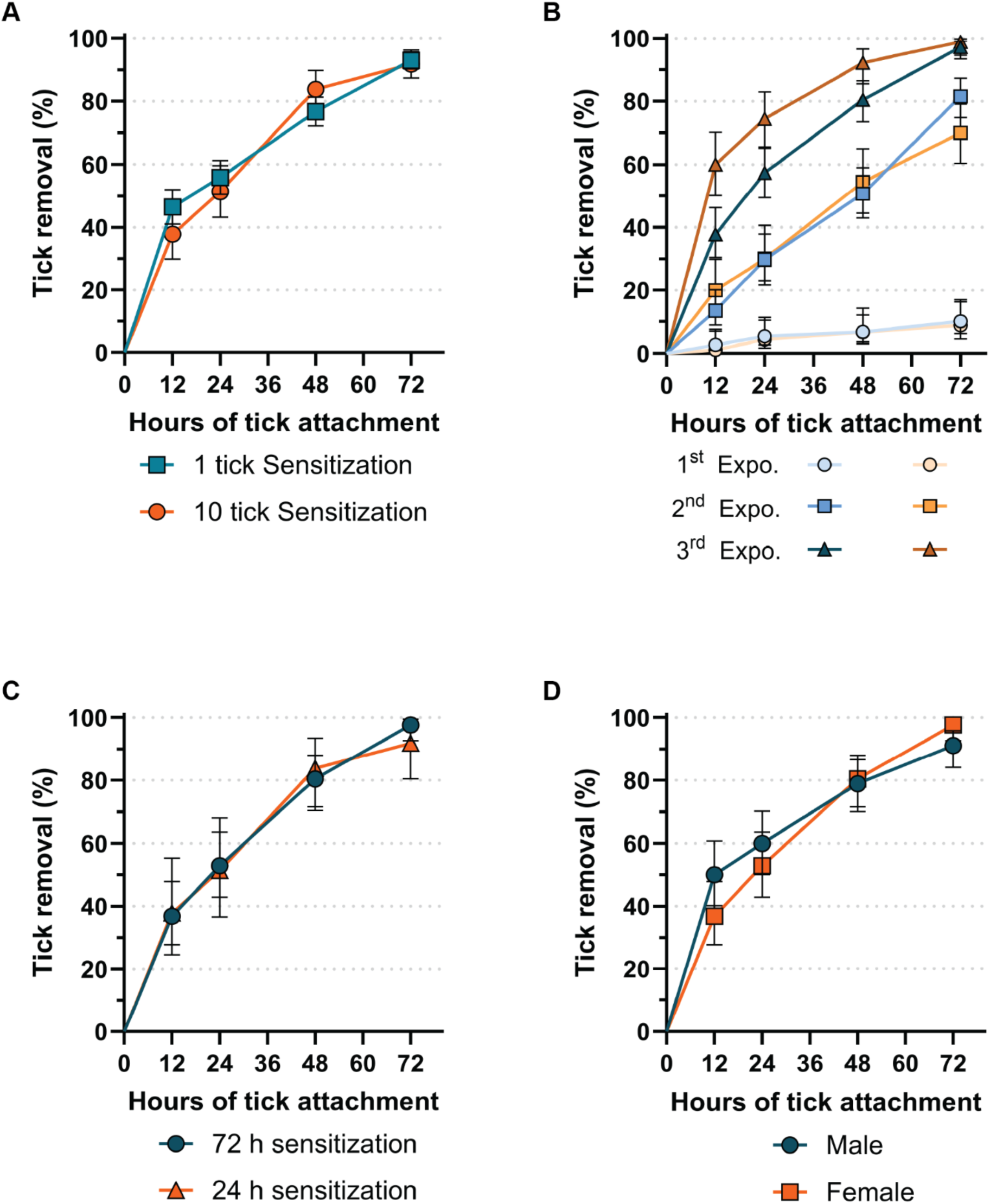
(**A-D**) Smoothed, inverted Kaplan-Meyer plot showing probability of tick removal (%). Analysis by Peto & Peto modification of the Gehan-Wilcoxon test. 95% CI shown. (**A**) 4^th^ and 3^rd^ weekly exposure of GPs sensitized with 1 and 10 ticks, respectively. p=0.782. (**B-D**) GPs were exposed to 15 ticks per exposure. (**B**) Tick removal success in three times sensitized GPs exposed weekly (blue shades) or every three weeks (orange shades). 1^st^ Exposure (Expo.): p=0.8787; 2^nd^ Expo.: p=0.8787; 3^rd^ Expo.: p=0.0006. (**C-D**) GPs were sensitized weekly. (**C**) Third exposure of GPs sensitized over 24 h or 72 h per exposure. p=0.8484. (**D**) Third exposure of male and female GPs. p=0.3898. For detailed statistics see supplementary report.

**Figure S8.**
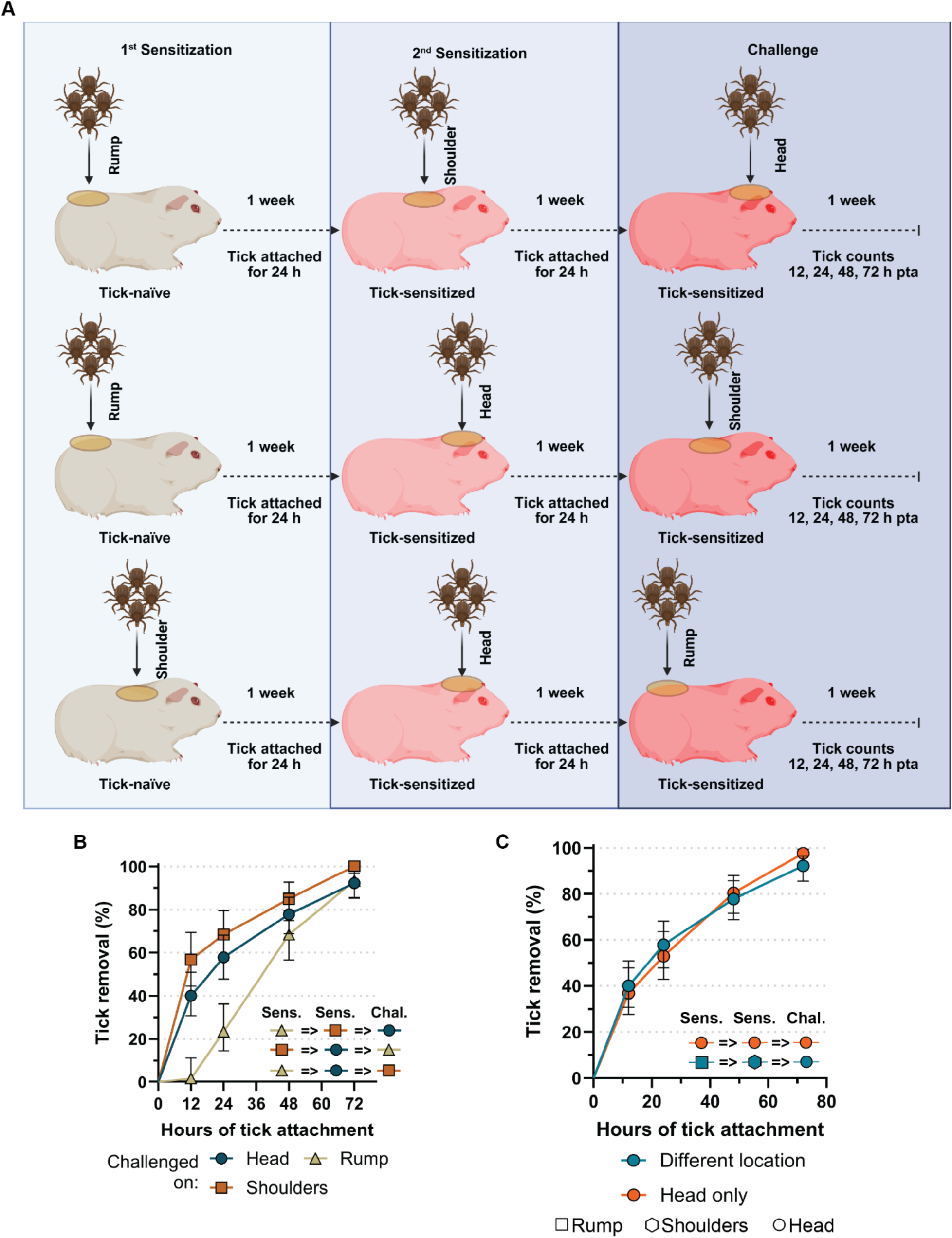
(**A**) Schematic of guinea pig (GP) sensitization and challenge depicting tick placement on different body locations (Created in BioRender: https://BioRender.com/y2smy29). (**B-C**) Smoothed, inverted Kaplan-Meyer plot showing probability of tick removal (%) in GPs sensitized weekly with 15 ticks. Analysis by Peto & Peto modification of the Gehan-Wilcoxon test. 95% CI is shown. (**B**) Shows results of tick challenge on different body location on GPs sensitized on other body locations than the challenge (see (**A**)). Head versus Rump: p<0.0001, Shoulders versus Rump: p<0.0001, Head versus Shoulders: p0.0483. (**C)** Tick removal success during a third exposure on the head in GP sensitized either on the same or different locations. p=0.9403. For more details see supplementary report.

**Fig. S9.**
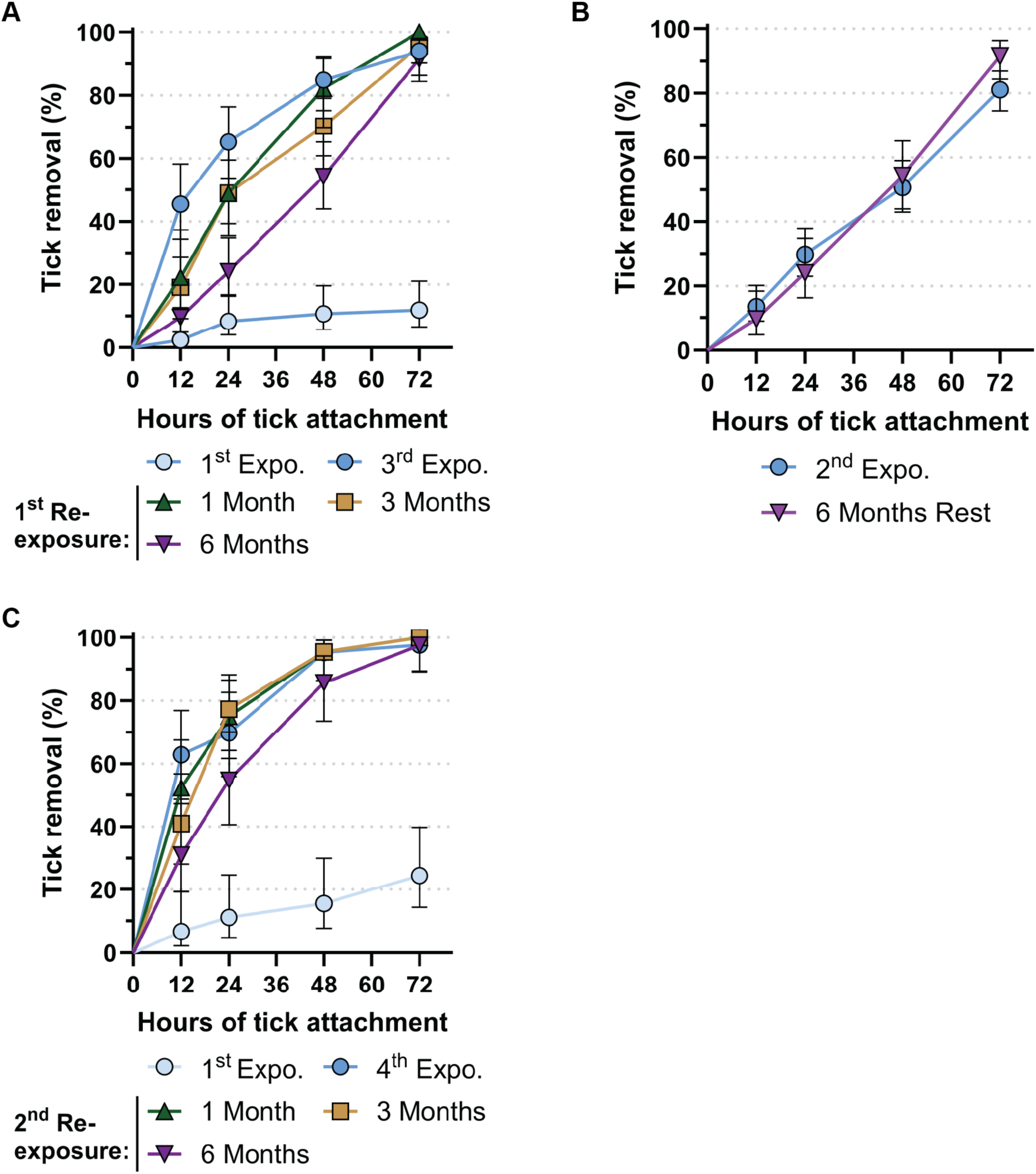
(**A-C**) Smoothed, inverted Kaplan-Meyer plot showing probability of tick removal (%). Analysis by Peto & Peto modification of the Gehan-Wilcoxon test. 95% CI is shown. **(A-C)** Guinea pigs (GPs) were exposed to 15 ticks. (**A**) Tick removal success in GPs after 1, 3, and 6 month(s) of rest (no tick exposure) compared to a first exposure of tick-naïve GPs or a third exposure of weekly sensitized GPs. Third exposure (expo.) versus 1 Month [rest] (p=0.0536) / versus 3 Months [rest] (p=0.0025) / versus 6 Months [rest] (p<0.0001). (**B**) Tick removal success of GPs during a second weekly exposure and after 6 months of no tick encounter. p=0.6281. (**C**) Tick removal success of the same GPs as in panel (**A**) during a subsequent tick-exposure a week later showing enhancement of tick removal success after long breaks from tick exposure. Forth exposure (expo.) versus 1 Month [rest] (p=0.6205) / versus 3 Months [rest] (p=0.3319) / versus 6 Months [rest] (p=0.03). For more details see supplementary report.

**Table S1.** Upregulated skin transcripts at the tick-bite site of tick-sensitized versus tick- tick-naive GP.

| Transcript name and link to Uniprot Mouse or Rat ortholog | NCBI Identifier and link | Description | Log(2) fold change | P-value | Function | Reference |
| --- | --- | --- | --- | --- | --- | --- |
| **Interleukins and other immune signaling products** |  |  |  |  |  |  |
| Il19 | 100715337 | interleukin 19 | 6.702564 | 5.25E-06 | This cytokine is found to be preferentially expressed in monocytes. It can bind the IL20 receptor leading to the activation of transcription 3 (STAT3). | (61) |
| Fgf23 | 100717518 | fibroblast growth factor 23 | 7.578738 | 0.000567 | Involved in positive regulation of B cell proliferation | (62) |
| Osm | 100724699 | oncostatin M | 4.402756 | 2.18E-09 | Oncostatin M (OSM) contributes to extracellular matrix remodeling, hematopoiesis, differentiation, inflammatory response and proliferation. | (63) |
| Il13 | 10072011<br>8 | interleukin 13 | 5.68252 | 0.009365 | This gene encodes an immunoregulatory cytokine produced primarily by activated Th2 cells. This cytokine is involved in several stages of B-cell maturation and differentiation. | (64) |
| Cd28 | 10071496<br>6 | CD28 molecule | 3.558919 | 0.000983 | Involved in immune response, T cell activation; T cell receptor signaling pathway; and in positive regulation of T cell proliferation; | (65) |
| Cd69 | 10072795<br>0 | CD69 molecule | 3.525725 | 5.66E-06 | Expression Cd69 is induced upon activation of T lymphocytes and may play a role in their proliferation. Furthermore, the protein may act to transmit signals in natural killer cells and platelets. | (66) |
| Il24 | 10071487<br>3 | interleukin 24 | 3.523712 | 0.002487 | IL-24 induces rapid activation of Stat-1 and Stat-3 transcription factors. | (67) |
| Fcrlb | 10071539<br>6 | Fc receptor like B | 3.512076 | 3.69E-05 | FCRL1-5 could regulate different features of B-cell evolution such as development, differentiation, activation, antibody secretion and isotype switching. | (68) |
| Ms4a7 | 10071948<br>5 | membrane spanning 4-domains A7 | 3.214635 | 0.000782 | This family member is associated with mature cellular function in the monocytic lineage, and it may be a component of a receptor complex involved in signal transduction. | (69) |
| Cd38 | 10602653<br>5 | ADP-ribosyl cyclase | 3.388717 | 0.001823 | Involved in positive regulation of B cell proliferation. | (70, 71) |
| Cd38 | 10072653<br>7 | CD38 molecule | 3.087273 | 2.44E-06 | Involved in positive regulation of B cell proliferation | (70, 71) |
| G3V7I1 | 10073536<br>8 | C-C motif chemokine 4 | 3.383439 | 0.000379 | Involved in the immune response. | (72) |
| Q5Y4N7 | 10072781<br>1 | adhesion G protein-coupled receptor E4P | 4.339531 | 0.004216 | Enables G protein-coupled receptor activity. | (73) |
| Fut7 | 10071738<br>5 | fucosyltransferase 7 | 3.82816 | 0.000981 | Involved in fucosylation. | (74) |
| **Cytotoxic T cells product** |  |  |  |  |  |  |
| Clec1b | 10072628<br>9 | C-type lectin domain | 3.592165 | 6.55E-09 | Natural killer cells express multiple calcium-dependent lectin-like receptors that either inhibit or activate cytotoxicity and cytokine secretion | (75) |
| GRAB | 10073618<br>0 | granzyme B | 6.598226 | 2.23E-06 | Granzyme B (granzyme 2, cytotoxic T-lymphocyte-associated serine esterase 1) | (76) |
| GRAC | 10071966<br>9 | granzyme C | 5.393386 | 0.000195 | Granzyme C (granzyme 2, cytotoxic T-lymphocyte-associated serine esterase 1) | (77) |
| GRAB-Like2 | 10072021<br>9 | granzyme B-like | 4.93077 | 6.42E-07 | Granzyme B (granzyme 2, - cytotoxic T-lymphocyte associated serine esterase 1) | (76) |
| **Chemotaxis** |  |  |  |  |  |  |
| LOC100734903 | 10073490<br>3 | C-C motif chemokine 3-like | 3.424106 | 0.000368 | Involved in eosinophil chemotaxis. | (78) |
| LOC100735187 | 10073518<br>7 | C-C motif chemokine 3-like | 3.132743 | 0.000621 | Involved in eosinophil chemotaxis. | (78) |
| LOC100731901 | 10073190<br>1 | alveolar macrophage chemotactic factor | 3.996885 | 1.19E-08 | Involved in immune response; involved in chemotaxis; and in defense response. | (79) |
| LOC100732171 | 10073217<br>1 | platelet basic protein-like | 5.654068 | 5.28E-05 | Involved in chemotaxis | (80) |
| **Proteases** |  |  |  |  |  |  |
| Matrix metalloproteinases |  |  |  |  |  |  |
| Mmp3 | 10072910<br>1 | stromelysin-1 | 3.872797 | 1.80E-11 | Involved in collagen catabolic process; and in extracellular matrix organization. | (81) |
| Mmp7 | Mmp4 | matrix metalloproteinase 7 | 4.247669 | 4.95E-07 | Involved in collagen catabolic process; and in extracellular matrix organization. | (82) |
| Mmp8 | 10072467<br>0 | matrix metalloproteinase 8, neutrophil collagenase | 3.120398 | 2.30E-05 | Involved in collagen catabolic process; and in extracellular matrix organization. | (83) |
| Mmp10 | 10072769<br>0 | matrix metalloproteinase 10 | 4.925036 | 1.64E-10 | Involved in collagen catabolic process; and in extracellular matrix organization. | (84) |
| Mmp13 | 10073175<br>6 | matrix metalloproteinase 13 | 3.183712 | 1.77E-05 | Involved in collagen catabolic process; involved in extracellular matrix organization. | (85) |
| Other peptidases |  |  |  |  |  |  |
| Kelz | 10072947<br>3 | Kelz metallo-endopeptidase | 4.564278 | 4.43E-05 | Hydrolyses bradykinin and neuropeptides. | (86, 87) |
| Prss29 | 1007243<br>32 | Serine protease 29-like | 6.95884<br>2 | 1.79E-05 | Involved in proteolysis. | (88) |
| **Prostanoid catabolism** |  |  |  |  |  |  |
| CP4FE | 100733735 | cytochrome P450 4F3-like | 5.74894 | 7.25E-05 | Cytochrome P450 4F3 (CYP4F3), originally identified as one of the leukotriene B4 ω-hydroxylases, participate in the metabolism of various endobiotics, as well as some xenobiotics. | (89) |
| **Innate immunity** |  |  |  |  |  |  |
| APOBE C-1 | 100730186 | APOBEC-1-like | 5.978293 | 0.007912 | Nuclear C to U RNA editing - many of these candidate RNAs are involved in cytokine signaling; anti-viral. | (90) |
| Oasl | 100734390 | 2'-5'-oligoadenylate synthetase like | 3.94653 | 4.14E-05 | Involved in defense response to virus; involved in negative regulation of viral genome replication; involved in regulation of ribonuclease activity | (91) |
| NGAL | 100722819 | neutrophil gelatinase-associated lipocalin-like | 6.307814 | 0.0051 | Enables iron ion binding activity. Involved in innate immune response, positive regulation of cold-induced thermogenesis; and siderophore transport. | (92) |
| Kunitz serine protease inhibitor |  |  |  |  |  |  |
| <u>Spint3</u> | <u>1017879</u><br><u>04</u> | Protease<br>inhibitors-like,<br>Kunitz type, 3 | 3.86132<br>8 | 3.76E-05 | Protease inhibition. | (93) |

